# Long noncoding RNA *MEG3* activates neuronal necroptosis in Alzheimer’s disease

**DOI:** 10.1101/2022.02.18.480849

**Authors:** Sriram Balusu, Katrien Horré, Nicola Thrupp, An Snellinx, Lutgarde Serneels, Iordana Chrysidou, Amaia M. Arranz, Annerieke Sierksma, Joel Simrén, Thomas K. Karikari, Henrik Zetterberg, Wei-Ting Chen, Dietmar Rudolf Thal, Evgenia Salta, Mark Fiers, Bart De Strooper

## Abstract

Neuronal cell loss is a defining feature of Alzheimer’s disease (AD), but it remains unclear how neurons die and how this relates to other defining characteristics of the disease^1^. Existing *in vivo* AD models only partially recapitulate the neuropathology of AD with very mild or no neuronal cell loss. Here we demonstrate that human neurons xenografted in mouse brain exposed to amyloid pathology develop sarkosyl-insoluble tau filaments, positive Gallyas silver staining, release phosphorylated tau (P-tau181) into the blood, and display considerable neuronal cell loss, providing a model for the induction of full Tau pathology by simple exposure to amyloid pathology in AD. The alterations are specific to human neurons and contrast with the mild effects exhibited in mouse neurons. A core transcriptional program in the human neurons is characterized by strong upregulation of *MEG3,* a neuron-specific long noncoding RNA*. MEG3* is also strongly upregulated in neurons from AD patients *in situ*. *MEG3* expression alone is sufficient to induce necroptosis in human neurons *in vitro*. Orally administered small molecule receptor-interacting protein (RIP) kinase -1 and -3 inhibitors rescued the neuronal cell loss in this novel AD model. Thus, xenografted human neurons are uniquely sensitive to amyloid pathology, recapitulate all the defining neuropathological features of AD, and ultimately die by necroptosis.

## Main text

The major question of how the defining hallmarks of Alzheimer’s disease (AD) amyloid-β (Aβ) plaques, neuronal tau tangles, granulovacuolar neurodegeneration and neuronal cell loss relate to each other has never been solved, largely owing to a lack of good models that encompass all neuropathological aspects of the disease. Aβ pathology strongly influences tau pathology kinetics in several mouse models, but only if tau pathology is artificially instigated by frontotemporal dementia (FTD) causing mutations or by injecting tau-seeds isolated from AD patient brains^2–4^. The fundamental unanswered question remains whether and how these protein accumulations cause neuronal loss in AD.

The many attempts to generate a complete model for AD have failed likely because of (unknown) human-specific features that escape modeling in rodents. We generated a much-improved mouse model for xenotransplantation of human neurons using a *Rag2^-/-^* (Rag2^tm1.1Cgn^) genetic background and a single *App^NL-G-F^* (App^tm3.1Tcs^/App^tm3.1Tcs^) knock-in gene to drive Aβ pathology, instead of the combined *APP-PS1* (Tg(Thy1-APP*Swe, Thy1-PSEN1*L166P)21Jckr) transgene overexpression used before^5^. Human stem cell-derived neuronal progenitor cells (NPCs) transplanted in the control *Rag2^-/-^* mice integrate well and develop over time dendritic spines, indicative of functionally active mature neurons (Extended Data Fig. 1a-b). Two months post-transplantation (2M PT), the xenografted neurons display characteristic mature neuronal (NEUN and MAP2) and cortical markers (CTIP2, SATB2, TBR1, and CUX2). Furthermore, unlike rodent neurons, human neurons display 3R and 4R tau splice forms at 6-months post-transplantation (6M PT) (Extended Data Fig. 1c). Compared to the previously used NOD-SCID animals^5^, *Rag2^-/-^* animals show a considerably increased life span (>18 months), allowing the study of healthy human neurons during brain aging.

**Figure 1:**
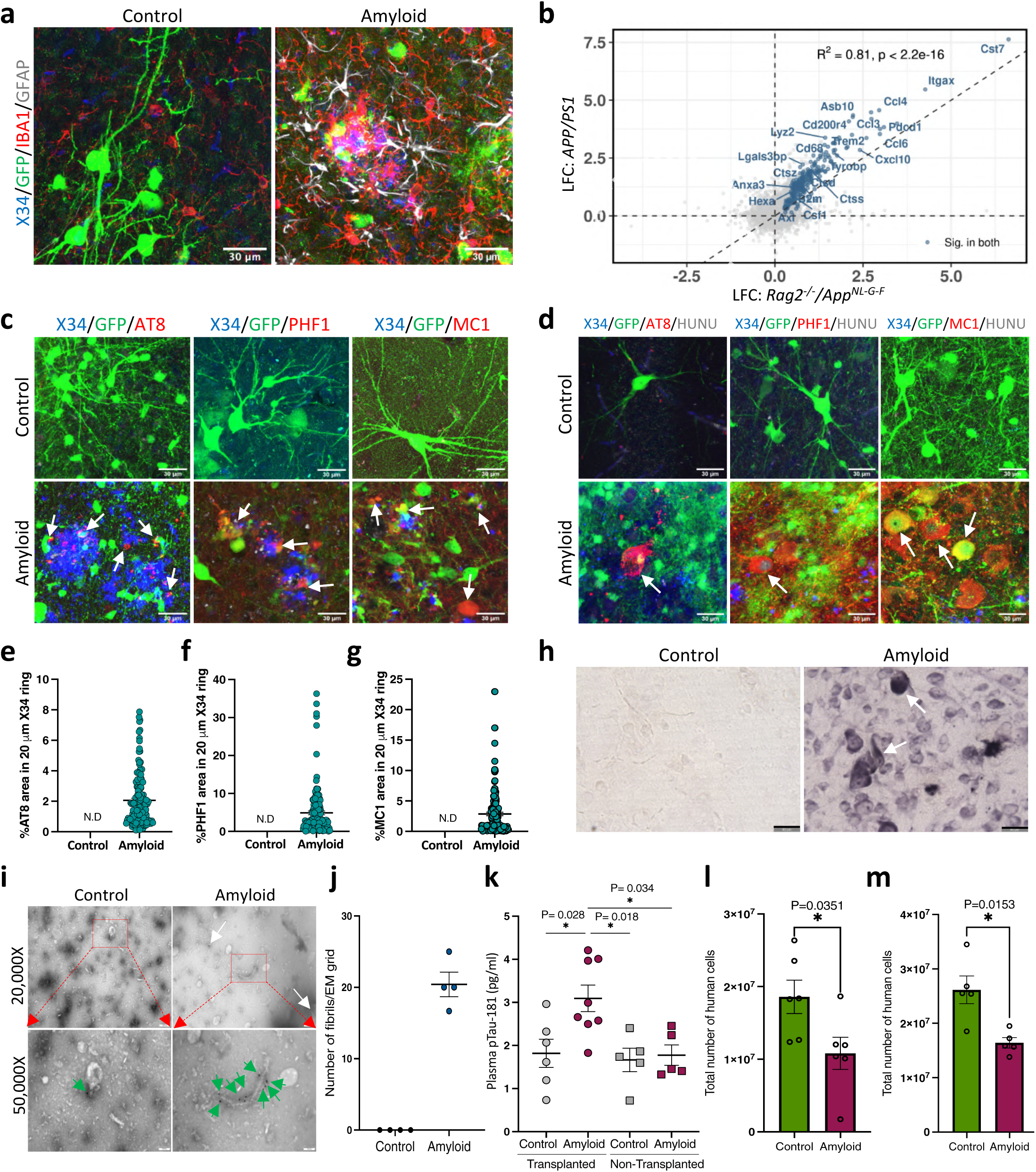
Amyloid plaque deposition is sufficient to induce pathological TAU in grafted human neurons but not in mouse neurons. **a,** Representative confocal images of grafted neurons of 18-months old control (left, n=4) and amyloid (right, n=4) mice. Grafted human neurons (GFP, green), amyloid (X34, blue), microglia (IBA1, red), astrocytes (GFAP, grey). Scale bar 30 µm. **b,** Scatter plot showing comparison of log fold changes (LFC) in *APP-PS1* mice at 10M and *Rag2^-/-^ /App^NL-G-F^* mice at 6M. Genes highlighted blue are significantly changed in both mouse models. The Pearson correlation (R^2^) of genes significantly changing in both models is 0.81. **c,** Representative confocal images showing neuritic plaque tau (NP-tau) in 18-months old control (n=4) and amyloid (n=4) mice. White arrows indicate X34 and NP-tau positive plaques. Amyloid plaques (X34, blue), grafted human neurons (GFP, green), AT8 (P-tau Ser202-Thr205, red), PHF1 (P-tau Ser396, red), and MC1 (pathological conformation, red). Scale bar 30 µM. **d,** Representative confocal images showing pathological tau accumulation in the neuronal somas in 18-months old control (n=4) and amyloid (n=4) mice. Some human neurons, mainly in amyloid mice, lost their immunoreactivity to the GFP antibody. Hence human nucleus (HUNU) antibody is also used to label the human cells. White arrows indicate P-tau or MC1 positive human neurons. Amyloid (X34, blue), grafted human neurons (GFP, green), AT8 (P-tau Ser202-Thr205, red), PHF1 (P-tau Ser396, red), MC1 (pathological conformation, red) and human nucleus (HUNU, grey). Scale bar 30 µM. **e,** Quantification of AT8 positive tau around Aβ plaque, within 20 microns diameter (n=4, >100 plaques/mice). ND, not detected. **f,** Quantification of PHF1 positive tau around Aβ plaque, within 20 microns diameter (n=4, >100 plaques/mice). ND, not detected. **g,** Quantification of MC1 positive tau around Aβ plaque, within 20 microns diameter. (n=4, >100 plaques/mice). ND, not detected. **h,** Representative light microscope Gallyas silver stain images from control (n=4) and amyloid (n=4) grafted animals. Scale bar 50 µm. **i,** Representative electron micrograph images of immunogold labelled sarkosyl insoluble tau fibrils isolated from 18-months old control (n=4) and amyloid (n=4) mice. Tau fibrils are indicated with white (top panel) or green (bottom panel) arrows. Top panel, 20,000x magnification (scale bar, 200 nm). Bottom panel, 50,000x magnification (scale bar, 100 nm). **j,** Quantification of the number of immunogold positive fibrils identified in the grafted control (n=4) and amyloid mice (n=4). **k,** Quantification of the plasma P-tau181 levels from 18-months old grafted (control n=6, amyloid n=8) and non-grafted (control n=5, amyloid n=5) mice. **l,** Quantitative representation of the human neurons at 6-months post-transplantation in control (n=6) and amyloid (n=6) mice. **m,** Quantitative representation of the human neurons at 18-months post-transplantation in control (n=5) and amyloid (n=6) mice. Values are presented as mean ± SEM. One-way ANOVA with Tukey’s post hoc test for multiple comparisons was used in (j), and Student’s t-test in (k, l) to measure the statistical significance.

We xenografted GFP-labelled H9-derived human cortical neuronal precursor cells (100,000 NPCs/mouse) into *Rag2^-/-^/App^NL-G-F^* (further referred to as amyloid mice) or *Rag2^-/-^/App^mm/mm^* (further referred to as control mice). β-sheet staining with dye X34 revealed robust plaque pathology in the amyloid mice (Fig. 1a & Extended Data Fig. 1d). RNA sequencing confirmed that the transcriptional profile of the human transplants at 2M in control and amyloid mice are very similar (Fig. 2a). Thus, human neurons integrate and differentiate similarly in control and amyloid mice, in line with the fact that amyloid plaque pathology appears only from 3M onwards in this model.

**Figure 2:**
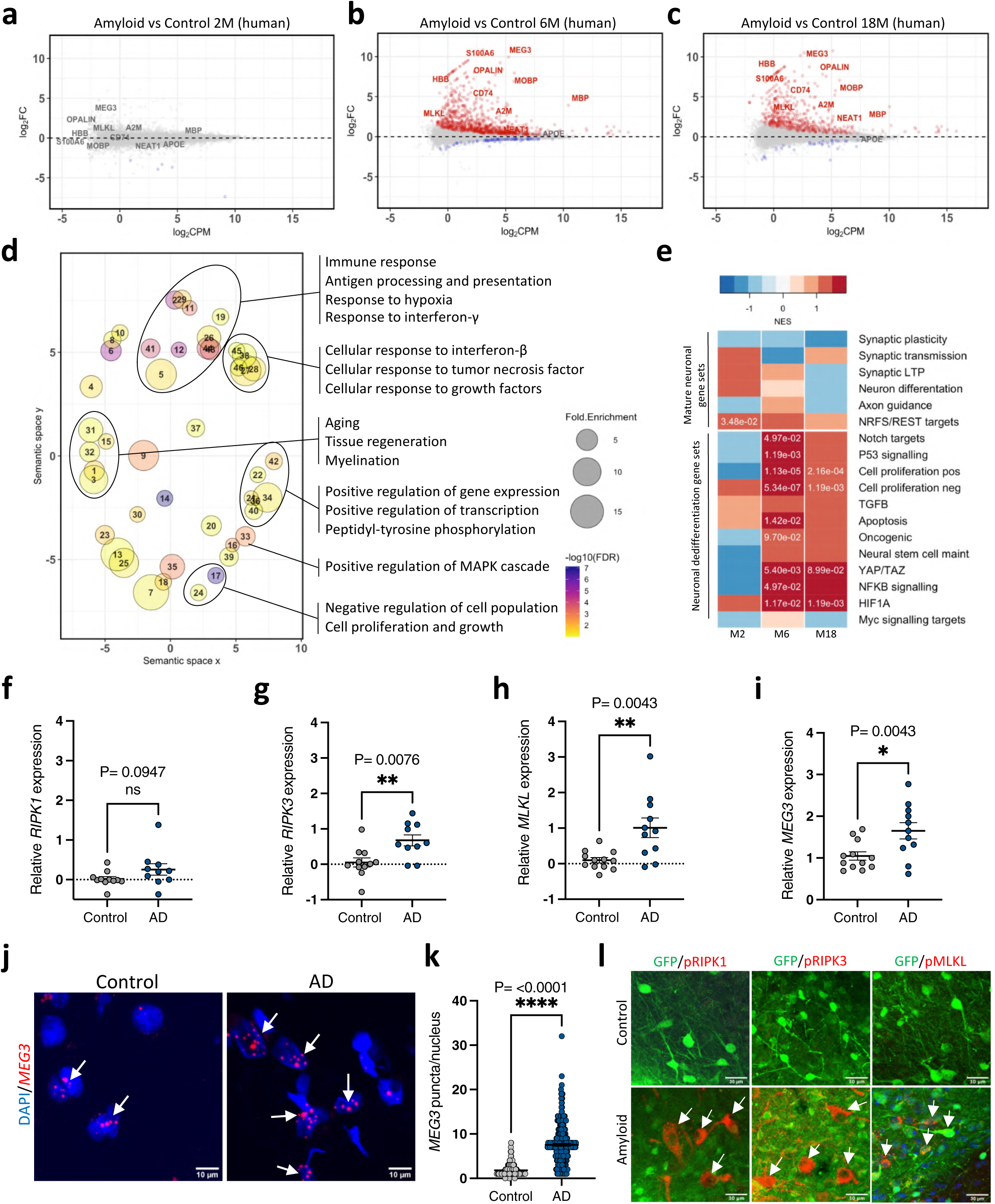
Transcriptional changes in xenografted neurons. Bland-Altman MA plot showing differential expression of genes from RNA sequencing of the human grafts isolated from randomized control and amyloid mice at **(a)** 2 months (control n=5, amyloid n=7), **(b)** 6 months (control n=5, amyloid n=5), and **(c)** 18months (control n=4, amyloid n=3) post-transplantation. Red, significantly upregulated genes. Blue, significantly downregulated genes (FDR < 0.05). FC=fold change, CPM=counts per million. Comparisons of logFC at month 6 and month 18 are shown in Extended data figure 5g. **d,** Gene Ontology (GO) analysis showing terms associated with genes upregulated in amyloid mice at 6M. Point size represents the fold enrichment of upregulated genes in the term, and color represents the –log_10_FDR. Only terms with FDR < 0.1 are shown. The x and y axes represent the ’semantic space’ - that is, how similar two terms are to each other (calculated by Revigo, PMID: 21789182). **e,** Heatmap of normalised enrichment scores (NES) in neuronal dedifferentiation gene sets ranked along the differentially expressed genes in the xenografts (amyloid vs control) shown in (Fig. 2a-c). Positive enrichments are shown in red, negative enrichments are shown in blue. Significant FDR values (P.adj < 0.01) are shown as numbers. **f,** Analysis of *RIPK1* gene expression using qPCR on AD (n=11) and control (n=10) post-mortem human brain samples. **g,** Analysis of *RIPK3* gene expression using qPCR on AD (n=11) and control (n=10) post-mortem human brain samples. **h,** Analysis of *MLKL* gene expression using qPCR on AD (n=11) and control (n=10) post-mortem human brain samples. **i,** Analysis of *MEG3* gene expression using qPCR on AD (n=11) and control (n=10) post-mortem human brain samples. **j,** Representative confocal images showing *MEG3* RNAScope in temporal gyrus of AD (n=3) and control (n=2) post-mortem human brain samples. Nucleus (DAPI, blue), *MEG3* (red). Scalebar 10 µM. **k,** Quantitative representation of the number of *MEG3* puncta per nucleus in control (>100 nuclei/sample, n=2), and AD (>100 nuclei/sample, n=3) post-mortem human brain samples. **l,** Representative confocal images showing the expression of activated necroptosis pathway markers in 18-months old human neurons in control (n=4) and amyloid (n=4) mice. Grafted human neurons (GFP, green), pRIPK1, pRIPK3 or pMLKL (red). White arrows indicate pRIPK1, pRIPK3, or pMLKL positive cells. Scalebar 30 µm.

Full blown amyloid plaque pathology is seen at 18M PT. At this late point in time, human neurons in the control brain appear overall healthy, display neuronal projections and dendritic spines intermingled with host microglial (IBA1) and astroglia (GFAP) cells (Extended Data Fig. 1d). In contrast, grafted neurons in the amyloid mice display severe dystrophic neurites near Aβ plaques, associated with microgliosis (6-7 IBA1-positive microglia per Aβ plaque) and astrogliosis (2-3 GFAP-positive astrocytes per plaque) (Fig. 1a, Extended Data Fig. 2a, b). The glial cell recruitment to plaques is very similar to non-grafted control animals (Extended Data Fig. 3a, b). Immunohistochemistry with phosphorylated tau (P-tau) antibodies, AT8 (P-tau Ser202, Thr205), PHF1 (P-tau Ser396), and MC1 (pathological conformational tau epitope) demonstrate considerable deposition of neuritic plaque tau (NP-tau)^2^ in the human neurons of the grafted amyloid mice (Fig. 1c, d) (Surface stained around Aβ plaques (20 µm ring) was ∼2% (AT8), ∼5% (PHF1), and ∼3% (MC1), Fig. 1e-g). Interestingly, both NP-tau- and AT8-positive neurons appeared as early as 6-months post-transplantation (Extended Data Fig.1e), indicating that amyloid deposition drives tau phosphorylation early on in this model. Significant tau pathology is not observed in mouse neurons in the same animal, nor in grafted control, nor in non-grafted amyloid mice (Fig. 1c, d, Extended Data Fig. 1e (top panel) & Extended Data Fig. 3c).

**Figure 3:**
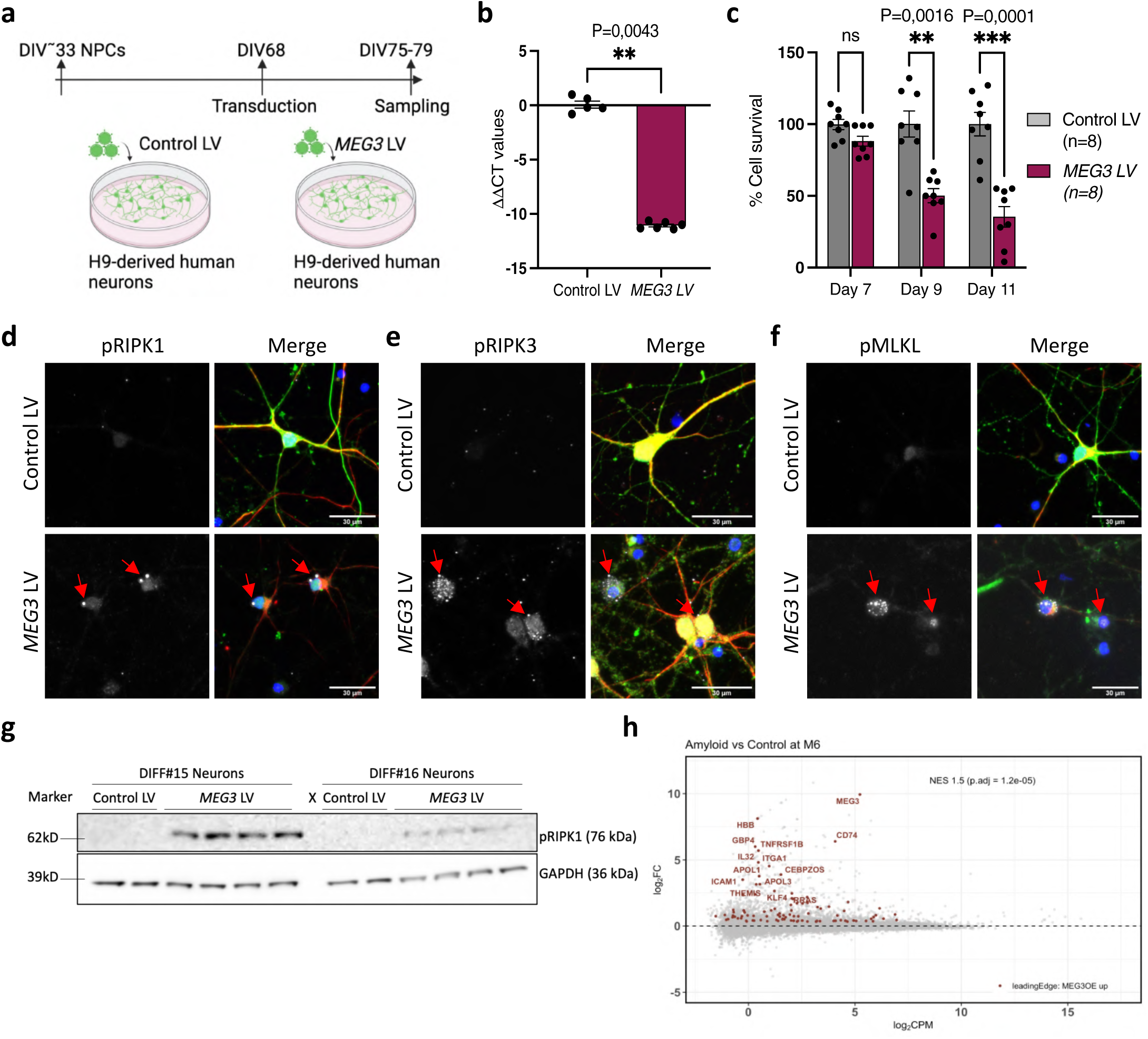
Long noncoding RNA *MEG3* induces necroptosis in human neurons. **a,** Schematic representation of the *MEG3* expression strategy using lentiviral vectors (LV) in H9- derived human neurons. **b,** Analysis of the *MEG3* expression in H9-derived neurons at DIV75, seven days post-transduction with either control LV (n=5) or *MEG3* LV (n=6). **c,** Analysis of neuronal cell survival using CellTiter-Glo reagent after transducing with either control LV (n=8) or *MEG3* LV (n=8) on day-7, day-9, and day-11 post-transduction. **d,** Representative confocal image showing activated necroptotic marker (pRIPK1, Ser 166) in H9- derived neurons at DIV75 transduced with either control LV (n=3) or the *MEG3* LV (n=3). Nucleus (DAPI, blue), pRIPK1 (grey, indicated with red arrows), transduced neurons (GFP, green), neurons (NF-H, red). Scalebar 30 µm. **e,** Representative confocal image showing activated necroptotic marker (pRIPK3, Ser 227) in H9- derived neurons at DIV75 transduced with either control LV (n=3) or the *MEG3* LV (n=3). Nucleus (DAPI, blue), pRIPK3 (grey, indicated with red arrows), transduced neurons (GFP, green), neurons (NF-H, red). Scalebar 30 µm. **f,** Representative confocal image showing activated necroptotic marker (pMLKL, Ser 358) in H9- derived neurons at DIV75 transduced with either control LV (n=3) or the *MEG3* LV (n=3). Nucleus (DAPI, blue), pMLKL (grey, indicated with red arrows), transduced neurons (GFP, green), neurons (NF-H, red). Scalebar 30 µm. **g,** Immunoblot analysis of pRIPK1 (Ser 166) levels. Proteins were extracted seven days post-transduction from H9-derived neurons at DIV75 transduced with either control LV (n=4) or *MEG3* LV (n=8). GAPDH is used as the loading control. Two independent differentiations (Diff). **h,** The same Bland-Altman MA plot showing differential expression of bulk RNA sequencing of human grafts from 6 months as in Fig. 2b. Genes highlighted in red are the leading-edge genes identified in a gene-set enrichment analysis (GSEA), taking the top 400 upregulated genes from the bulk sequencing of the primary neurons expressing *MEG3* (Extended Data Fig. 10) and plotting them against the amyloid vs control fold changes of the human grafts (FDR<0.05). FC=fold change, CPM=counts per million.

Pathological, β-sheet X-34-positive tau-reactivity is seen in the xenografted human neurons (Extended Data Fig. 2c). Furthermore, Gallyas silver staining and tau-immunogold labeling of PHF fibril-like structures extracted with sarkosyl from grafted neurons in the amyloid mice (Fig. 1h, i) confirmed the progression of P-tau into pathological states (> 20 tau-fibrils/EM grid (control n=4, amyloid n=4) (Fig. 1j). Such pathology is largely absent in grafted control or non-grafted amyloid mice (Fig. 1h, i & Extended Data Fig. 4a-c). Finally, and clinically relevant, we found that P-tau181^6, 7^ is significantly increased in the plasma of the grafted amyloid mice but not in control grafted or non-grafted mice (Fig. 1k), reflecting an increased secretion of soluble P-tau species from neurons into the bloodstream in response to amyloid pathology, mimicking what is observed in humans with AD.

**Figure 4:**
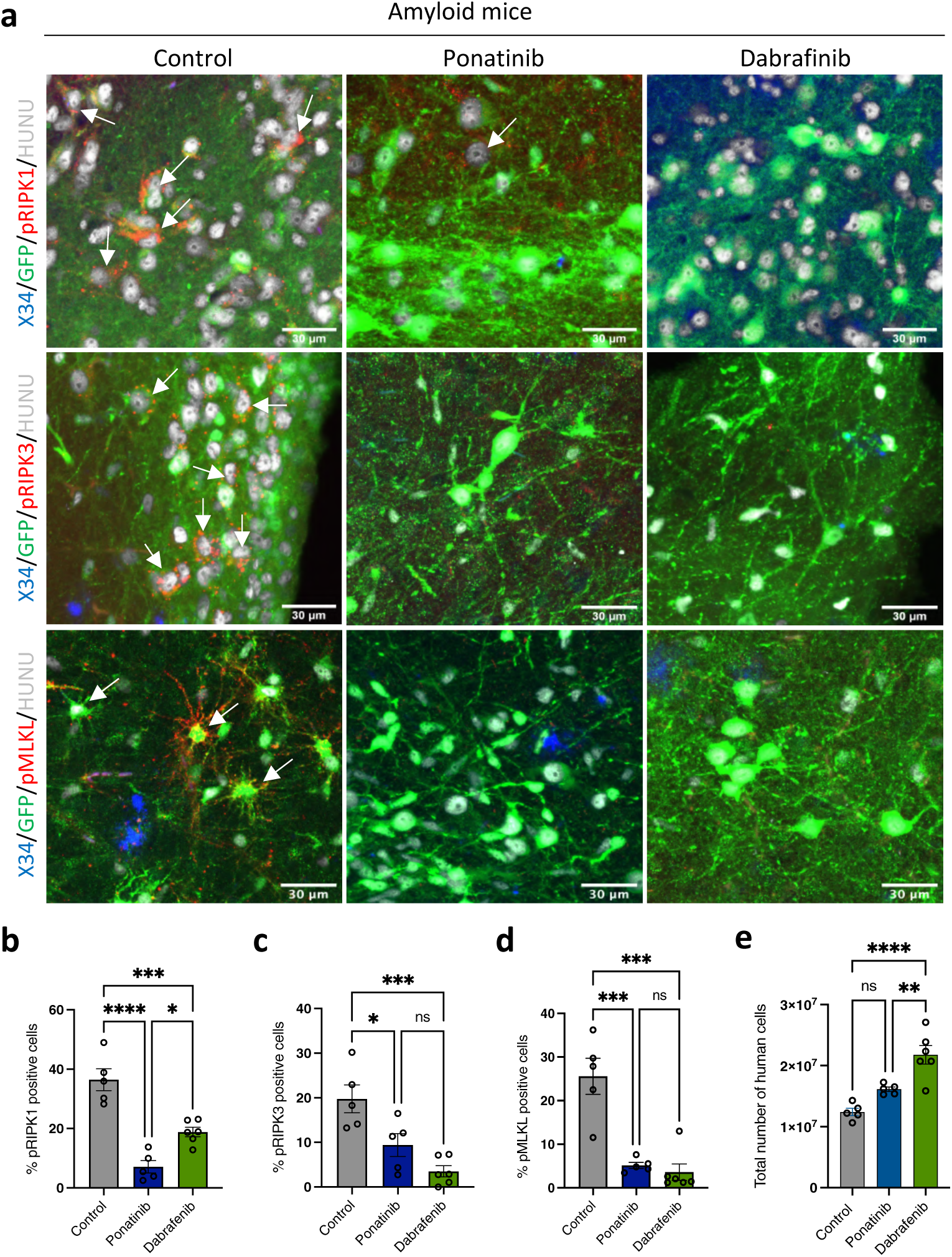
Inhibition of necroptosis prevents human neuronal cell loss *in vivo*. **a,** Representative confocal images showing the expression of the activated necroptotic markers pRIPK1, pRIPK3, and pMLKL in amyloid animals (n=5), amyloid animals treated with either ponatinib (n=5) or dabrafenib (n=6). Amyloid (X34, blue), human neurons (GFP, green), pRIPK1, pRIPK3, and pMLKL (red), human nuclei (HUNU, grey). White arrows indicate necroptosis marker positive neurons. Scale bar 30 µm. **b,** Quantitative representation of the percent number of human cells showing immunoreactivity to pRIPK1 in control (n=5), ponatinib (n=5), and dabrafenib (n=6) treated animals. **c,** Quantitative representation of the percent number of human cells showing immunoreactivity to pRIPK3 in control (n=5), ponatinib (n=5), and dabrafenib (n=6) treated animals. **d,** Quantitative representation of the percent number of human cells showing immunoreactivity to pMLKL in control (n=5), ponatinib (n=5), and dabrafenib (n=6) treated animals. **e,** Quantitative estimation of the number of human cells present in control (n=5), ponatinib (n=5), and dabrafenib (n=6) treated grafted mice at 6 months post-transplantation.

It is possible to quantify accurately up to 0.1 ng of human DNA in a mixture with 100 ng mouse genomic DNA using qPCR^8^ (Extended Data Fig. 6d). Using this assay, we estimate that up to 50% fewer human neurons were present in the amyloid animals compared to control animals (Fig. 1l, m) at 18M post transplantation (Extended Data Fig. 6e). The number of mouse neurons was also not significantly affected in the non-grafted control and amyloid mice (Extended Data Fig. 3e-g). Thus, human neurons, but not mouse neurons, develop robust neuropathological AD upon exposure to amyloid plaques *in vivo* and degenerate much like their counterparts in the human brain by an unknown death process.

Previous work has shown (opposing) effects of *Rag2^-/-^* immunodeficiency on amyloid plaque pathology and cellular responses^9, 10^. Comparison of soluble Aβ (Aβ40 & Aβ42) and guanidine-extractable Aβ (Aβ40 & Aβ42) revealed no significant differences between *Rag2^-/-^/App^NL-G-F^* and *App^NL-G-F^* mice (Extended Data Fig. 3h-k). Bulk RNA sequencing indicated that main aspects of the innate microglia and astroglial cellular transcriptomic responses to amyloid plaques are maintained in the *Rag2^-/-^/App^NL-G-F^* immune-deficient mouse compared to a previously characterized non-immune suppressed amyloid mouse (*APP/PS1*)^11, 12^ (Fig. 1b) (R^2^=0.81 for differentially expressed genes). Thus, while our model does not have an adaptive immune system, key features of the neuropathology of AD such as amyloid accumulation, glial responses, tau aggregation, tau phosphorylation, and neuronal cell loss, are reproduced in transplanted human neurons by simple exposure to amyloid pathology.

### Transcriptional changes in xenografted neurons reveal induction of necroptosis

We isolated the transplanted neuron-positive brain regions guided by the green fluorescent protein (GFP) marker from mice at 2-, 6-, and 18-months post-transplantation and extracted total RNA for sequencing (mean human reads ∼13.3 million) (Extended Data Fig. 5a, b). Mapping to mouse and human reference databases generated species-specific datasets. As expected, the mouse samples clustered according to genotype and age (Extended Data Fig. 5c, d). Moreover, upregulation of the mouse genes *Axl, B2m, Cst7, Ctss, Itgax, Trem2, Tyrobp, Lyz2, Ccl3,* and *Ccl4* confirms microglial activation when exposed to amyloid in the *Rag2^-/-^/App^NL-G-F^* mice (Fig. 1b, Extended Data Fig. 7a-c and Supplementary Table 2)^12^. Differential expression (DE) analysis of the human grafts at two months showed only a few significantly downregulated DE genes, of unknown relevance (Fig. 2a, Supplementary Table 1). In contrast, DE of grafts in amyloid and control conditions at 6M and 18M revealed 916 up and 73 down-regulated genes, and 533 up and 34 down-regulated genes, respectively (Fig. 2b, c and Supplementary Table 1). Gene alterations of interest include *CD74*, a marker for tau tangle-containing cells^13^; *HBB*, a cortical pyramidal neuronal marker^14–16^ and *A2M*, which is associated with neuritic plaques in AD human brains^17^ (Extended Data Fig. 5e). In addition to the neuronal signature, in line with our previous findings^18^, we also observe astroglial and oligodendrocytic genes, including *MOBP, MBP, OPALIN, S100A6, APOE,* among others (Extended Data Fig. 5f). The overall expression profiles of neurons exposed to amyloid plaques at 6M and 18M show strong correlations (R2 = 0.88 for genes differentially expressed at one or both time points), suggesting that the pathological cell state of neurons at 6M and at 18M are similar (Extended Data Fig. 5g).

We investigated whether the gene signatures of the human neurons exposed to amyloid in xenografted mice are overlapping with transcriptional changes observed in human AD brains (Supplementary Table 5). We found a remarkable enrichment using gene set enrichment analysis (GSEA) between previously published AD datasets, including the ROSMAP cohort^19^ and our data in transplanted neurons at 6M and 18M PT (P.adj <0.05) but not at 2M (Extended Data Fig. 8a, b). Furthermore, GSEA also revealed a striking enrichment in upregulated genes from transplanted neurons and neurons directly reprogrammed from fibroblasts of AD patients (M6: P.adj=1.64e-05, M18: P.adj=2.19e-04). These data confirm that our transplanted neurons capture AD-relevant transcriptional signatures^20^.

Functional gene ontology (GO) enrichment analysis of significantly upregulated genes (P.adj <0.05) at 6M using DAVID and REVIGO covered semantic space around positive regulation of transcription, protein phosphorylation, positive regulation of MAPK cascade, inflammatory responses, including TNF and interferon signaling, cell proliferation, aging, tissue regeneration, and myelination. (Fig. 2d and Supplementary Table 3). It is well established that neurons in AD display signatures of hypo-maturity, dedifferentiation, and cell cycle re-entry^20–23^. Using previously published datasets (Supplementary Table 4) and Gene set enrichment analysis (GSEA), we find that cellular signatures of hypo-maturity, dedifferentiation, P53, HIF*α* signaling, NF-kB, Myc, TGFβ-signaling, regulation of cell cycle (positive and negative), and cell-cycle re-entry are highly enriched in the 6M and 18M grafts but not at 2M (Fig. 2e). Conversely, mature neuronal marker pathways, such as synaptic plasticity, synaptic transmission, long-term potentiation, and axon guidance were not significantly enriched (Fig. 2e).

How neurons die in AD remains a highly controversial discussion^24, 25^. As shown in Fig. 1k-l, about ∼50% of xenografted human neurons are lost in the amyloid mice. We did not observe any significant alterations in the expression of genes associated with cell death mechanisms such as apoptosis or ferroptosis, but found significant upregulation of *MLKL*, the gene encoding the executor protein of necroptosis (Fig. 2b, c, Extended Data Fig. 2e & Extended Data Fig. 6a-c). This is not observed in the transcriptome of the host tissue (Extended Data Fig.7a-c). We used pRIPK1 (Ser 166), pRIPK3 (Ser 227), and pMLKL (Ser 358) specific antibodies to stain brain tissue from 18M grafted and non-grafted mice. This resulted in intense punctuate staining with a vesicular pattern in the soma of the neurons in grafted amyloid mice. These vesicular structures co-stained with casein kinase 1 delta (CK1*δ*), a marker for granulovacuolar degeneration (Fig. 2l and Extended Data Fig.6f). Non-grafted control and amyloid animals did not display these pathologies (Extended Data Fig. 3d). We validated this observation by qPCR on RNA extracted from the temporal gyrus of AD and age-matched control brain samples, which revealed significant upregulation of *MLKL* and *RIPK3*, an upstream kinase in the necroptosis pathway in the human brain (Fig. 2f-h). Based on initial observations made in the xenograft model described here, we have already extensively confirmed the presence of necroptosis markers in granulovacuolar degeneration in neurons of AD patients in a previous publication^26^. Thus, our model has predictive value for the neuropathology of AD.

### *MEG3* modulates the neuronal necroptosis pathway

We identified 36 up and 21 down-regulated long noncoding RNAs (lncRNA) in the grafts of 6-month old animals (Supplementary Table 1). Some of these non-coding RNAs (e.g., *NEAT1*) have been implicated in AD^27^, but overall it is unclear whether and how they contribute to pathogenesis. In our data, the long non-coding RNA *MEG3* (Maternally Expressed 3) was the most strongly (∼10 fold) upregulated gene in human neurons exposed to amyloid pathology (Fig. 2b-c & Extended Data Fig. 2d). *MEG3* has been linked to cell death pathways^28^ via p53^29^, is involved in the TGFβ pathway^30^, and has been associated with Huntington’s disease^31^. In a cerebral ischemia-reperfusion injury model, repression of *MEG3* has beneficial effects^32^.

While *MEG3* has not been previously implicated in AD, we found upregulation of *MEG3* in AD patient brains in a single nucleus transcriptomic database^33^. We confirmed a 2-3-fold upregulation of *MEG3* in RNA extracted from the temporal gyrus of human AD brains (Supplementary Table 8) using qPCR (Fig. 2i). *MEG3* in situ hybridization combined with cell-specific immunohistochemical markers NeuN (neurons), IBA1 (microglia), and GFAP (astrocytes) demonstrated its exclusive expression in the nucleus of neurons (Extended Data Fig. 9a, b) and its strong enrichment in AD brains (2-3 puncta/nucleus in control and 8-10 puncta/nucleus in AD brain (P= <0.0001), (Fig. 2j, k). In AD, *MEG3*-expressing neuronal nuclei appeared blotched with reduced DAPI intensity, and *MEG3-*positive neurons also displayed high levels of the necroptosis marker pMLKL (Extended Data Fig. 9c).

*MEG3* has 12 known transcriptional variants in humans^34^. We performed Sanger sequencing on brain cDNA and found that *MEG3* transcript variant-1 (accession number NR_002766.2) is most abundantly expressed in the human adult brain. A lentivirus was used to express *MEG3* V1 in H9-derived mature cortical neurons (Fig. 3a, b). Expression of *MEG3* in the neurons resulted in a strong reduction of cell viability at 9 days after transduction compared with control-transduced human neurons (Fig. 3c). Immunohistochemistry analysis revealed the presence of activated necroptotic markers, pRIPK1 (S166), pRIPK3 (S227), and pMLKL (S358) (Fig. 3d-f). Necroptosis-positive neurons displayed a reduced amount of cytoskeletal filament neurofilament-H (NF-H). Western blot analysis confirmed increased pRIPK1 (S166) levels (Fig. 3g), showing that the neurons were dying from necroptosis.

Next, to understand to what extent the *MEG3* expression contributes to transcriptional changes in the xenografted neurons, we performed RNA sequencing of *MEG3* transduced neurons 7-days post-transduction. GSEA of upregulated genes from *MEG3* expression *in vitro* against 6M xenografted neurons revealed a significant enrichment (NES=1.5, P.adj=1.2e-05) (Fig. 3h and Supplementary Table 7), suggesting that a subset of transcriptional changes in the xenografted neurons might be secondary to the upregulation of *MEG3*. However, DE analysis of *MEG3* transduced neurons revealed no changes in necroptosis genes (Extended Data Fig. 10a-c). DAVID Gene Ontology (GO) analysis of leading edge genes from GSEA on upregulated genes from 6M transplanted human neurons and *in vitro MEG3* expressing neurons revealed signatures of NF-kB and TNF signaling which have been related to necroptosis induction^35^. In addition, signatures of lipid transport, interferon-*γ* signaling, positive regulation of peptidyl-tyrosine phosphorylation, and apolipoprotein-L signaling are observed (Fig. 10d and Supplementary Table 7).

### Pharmacological inhibition of RIPK1 and RIPK3 blocks neuronal loss in xenografted mice

It remains unknown whether the neurons that degenerate during AD or in our neuronal transplants are all dying by necroptosis or whether necroptosis markers are limited to the neurons that are remaining at the stage of pathological analysis. We thus treated xenografted mice (3 groups of n=5) from 2M until 6M post-transplantation with the orally available necroptosis kinase inhibitors, ponatinib (30 mg/kg), and dabrafenib (50 mg/kg). Ponatinib inhibits RIPK1^36^ and RIPK3^37^ and is an FDA-approved drug for the treatment of acute lymphoid leukemia (ALL) and chronic myeloid leukemia (CML). Dabrafenib is a specific inhibitor of RIPK3^38^. Immunostaining at six months revealed a striking reduction in the levels of pRIPK1, pRIPK3, and pMLKL in both treatment groups compared to untreated mice (Fig. 4a-d). Excitingly, the qPCR assay analysis of neuronal cell numbers revealed a significant increase in neurons in the treated groups up to levels seen in the grafted control mice (without amyloid) (Fig. 4e and Fig. 1).

## Conclusion

Our results demonstrate that amyloid pathology is sufficient to induce full neuropathology of AD in non-genetically manipulated human neurons, including dystrophic neurites, tau fibrils, granulovacuolar degeneration, cell loss, and the appearance of the P-tau181 biomarker in the plasma of the grafted mice. One of the major unsolved questions in AD research is how neurons die. Some reports have proposed apoptotic mechanisms, but it has never been demonstrated convincingly that this drives neuronal cell loss in the disease^39^. While previous publications suggest activated necroptosis in the AD brain^26, 40^, descriptive studies cannot prove what happened in the neurons that were already lost. We demonstrate here that a large part of the degenerating neurons in the xenograft model can be rescued by treatment with clinically relevant necroptosis inhibitors^38^. Thus, we suggest that neuronal death in AD is largely driven by necroptosis, linking the loss of neurons to inflammatory processes that are upstream of this well-studied death pathway^12, 41, 42^. Therefore, therapies that prevent neuronal cell loss, in combination with more mainstream Aβ and tau targeted interventions, might be useful additions to the current efforts to develop disease-modifying strategies for AD^43–45^. Our data suggest that necroptosis is downstream of the accumulation of pathological tau and is induced by the upregulation of the non-coding RNA *MEG3* possibly via TNF-inflammatory pathway signaling. Long non-coding RNAs (lncRNAs) are important regulators of gene expression and influence a variety of biological processes, including brain aging and neurodegenerative disease^46^. The large number of non-coding RNAs that are differentially expressed in our novel AD model warrant further investigation.

It is deeply intriguing that human transplanted neurons display an AD phenotype, while mouse neurons interspersed within the graft do not display such signs. While human-specific features might define the sensitivity of the neurons to amyloid pathology, there is an attractive alternative hypothesis that gains some support from our transcriptional analysis showing signatures of downregulation of mature neuronal properties and upregulation of immature signalling pathways. This was also observed in a recent study that analyzed the transcriptional profiles in neurons directly derived from fibroblasts of AD patients (iNs)^20^. The neurons in that study did not show tangles, amyloid, or necroptosis markers typical of AD, and it was not clear how those observations relate to the classical hallmarks of AD. Our data suggest alternatively, that the relative immature profile of the xenografted human neurons in our model makes them sensitive to amyloid pathology, resulting in tau pathology and other hallmarks of AD. Paradoxically, we might have mimicked in our model the reactivation of immature and progenitor-like signaling pathways or DNA repair programs, also observed in neurons from patients at risk of AD^20, 47^, pointing to this age associated hypo-mature neuronal identity as an important factor in the development of full blown AD pathology.

## Acknowledgments

We thank Véronique Hendrickx and Jonas Verwaeren for animal husbandry. Sebastian Munck, Nikky Corthout, Axelle Kerstens, and Abril Escamilla Yala for their assistance in imaging (LiMoNe, VIB). Confocal microscope equipment was acquired through a Hercules Type 1 AKUL/09/037 to prof Wim Annaert. Dr. Pieter Baatsen, EM-platform of the VIB Bioimaging Core at KULeuven, for his assistance in electron microscopy imaging. *App^NL-G-F^* mice were kindly provided by Takaomi Saido (RIKEN Brain Science Institute, Japan).

SB is a Fonds voor Wetenschappelijk Onderzoek– Vlaanderen (FWO) senior postdoctoral scholar (12P5922N). AMA is a Ramon y Cajal Scholar supported by the Spanish Ministry of Science MCIN/AEI and FSE invierte en tu futuro (RYC2020-029494-I, RTI2018-101850-A-I00), and the Alzheimer’s Association US (AARG-21-85038). ES receives funding from Alzheimer’s Nederland and Health Holland. JS is supported by research grants from Stiftelsen för Gamla tjänarinnor, Demensfonden and Stohnes stiftelse. TKK is funded by the Swedish Research Council (Vetenskåpradet; #2021-03244), the Alzheimer’s Association (#AARF-21-850325), the BrightFocus Foundation (#A2020812F), the International Society for Neurochemistry’s Career Development Grant, the Swedish Alzheimer Foundation (Alzheimerfonden; #AF-930627), the Swedish Brain Foundation (Hjärnfonden; #FO2020-0240), the Swedish Dementia Foundation (Demensförbundet), the Swedish Parkinson Foundation (Parkinsonfonden), Gamla Tjänarinnor Foundation, the Aina (Ann) Wallströms and Mary-Ann Sjöbloms Foundation, the Agneta Prytz-Folkes & Gösta Folkes Foundation (#2020-00124), the Gun and Bertil Stohnes Foundation, and the Anna Lisa and Brother Björnsson’s Foundation. HZ is a Wallenberg Scholar supported by grants from the Swedish Research Council (#2018-02532), the European Research Council (#681712), Swedish State Support for Clinical Research (#ALFGBG-720931), the Alzheimer Drug Discovery Foundation (ADDF), USA (#201809-2016862), the AD Strategic Fund and the Alzheimer’s Association (#ADSF-21-831376-C, #ADSF-21-831381-C and #ADSF-21-831377-C), the Olav Thon Foundation, the Erling-Persson Family Foundation, Stiftelsen för Gamla Tjänarinnor, Hjärnfonden, Sweden (#FO2019-0228), the European Union’s Horizon 2020 research and innovation programme under the Marie Skłodowska-Curie grant agreement No 860197 (MIRIADE), European Union Joint Program for Neurodegenerative Disorders (JPND2021-00694), and the UK Dementia Research Institute at UCL. DRT receives funding from FWO (G0F8516N, G065721N) and Stichting Alzheimer Onderzoek (SAO-FRA 2020/017). The BDS laboratory is supported by a European Research Council (ERC) grant CELLPHASE_AD834682 (EU), FWO, KU Leuven, VIB, Stichting Alzheimer Onderzoek, Belgium (SAO), the UCB grant from the Elisabeth Foundation, a Methusalem grant from KU Leuven and the Flemish Government and Dementia Research Institute - MRC (UK). B.D.S. is the Bax-Vanluffelen Chair for Alzheimer’s Disease and is supported by the Opening the Future campaign and Mission Lucidity of KUL, Leuven University.

## Authors’ contribution

SB, BDS, conceptualized research; SB, KH, NT, AS, LS, IC, AAM, JS, TKK, HZ, WT-C, DRT, ES, MF designed methodology. SB, KH, AS, IC, AAM, AS, JS, WT-C, ES performed experiments. LS generated mouse strains. JS, TKK, HZ performed P-Tau181. MF, NT performed Bioinformatic analysis. SB and BDS wrote the manuscript with input from all authors. All authors reviewed and approved the manuscript.

## Materials and methods

### Generation of immunodeficient transgenic amyloid mice

*Rag2^-/-^* (*Rag2^tm1.1Cgn^*) and *App^NL-G-F^* (*App^tm3.1Tcs^*) mice in a C57BL/6 background were housed in our specific pathogen-free (SPF) animal facility. Homozygous *Rag2^-/-^* single knock-out mice and homozygous *App^NL-G-F^* transgenic knock-in mice were crossed to obtain the *Rag2^-/-^*/*App^NL-G-F^* genotype. Isogenic *Rag2^-/-^*/*App^mm/mm^* (*mm*, Mus *musculus*) containing the *App* wild type allele were used as control mice. Homozygous colonies were maintained. Crosses were conducted within the same genotype to obtain mice for grafting. Mice were randomized, and both sexes were used for the experiments. Mice were housed in groups of four per cage with ad libitum access to food and water and a 14-hour light/10hour dark cycle at 21 °C. For necroptosis inhibition, ponatinib (30 mg/kg) and dabrafenib (50 mg/kg) were mixed in the mouse food, which was replaced every three days during the treatment period. All rodent experiments were approved by the ethics committee of KU Leuven and were executed in compliance with the ethical regulation for animal research.

### Pluripotent stem cell culture and neuronal differentiation

All the experiments were performed using the established H9 stem cell line (WAe009-A; https://hpscreg.eu/cellline/WAe009-A), stably expressing GFP under the control of chicken b-actin promoter (CAG promoter). Routine culturing and maintenance of the stem cells was performed on Matrigel coated surface using E8-flex growth media (Thermo; #A2858501). Cells were maintained in a humidified chamber at 37°C with 5% CO2. Once the cells were confluent, they were passaged using 0.5 mM EDTA. The protocol described by Shi et al., 2012^1^, with minor modifications, was used to differentiate stem cells into cortical neurons. On day in vitro (DIV) -2 of neural induction, cells were enzymatically dissociated with StemPro™ Accutase™ Cell Dissociation Reagent (Sigma, # A1110501). Single cells were plated (2 x 10^6^/well) in Matrigel-coated 6-well plate in mTeSR1 plus growth medium (STEMCELL Technologies), supplemented with 10 mM Y-27632 ROCK inhibitor (Calbiochem, CA, USA). For neural induction, the culture medium was replaced with a neural maintenance medium (NMM) supplemented with 1 µM StemMACS™ LDN-193189 (Miltenyi Biotec, #130-106-540) and 10 µM SB431542 (STEMCELL Technologies, #72234) every day until DIV12. From DIV12 to DIV24, neuroepithelium was passaged three times using mechanically dissociation, to enrich for neural rosettes. Around DIV27, neural rosettes were dissociated into single cells using the StemPro™ Accutase™ Cell Dissociation Reagent. DIV30 cells were frozen as progenitors (NPCs) for future use.

### Neural progenitor cells (NPCs) culturing for grafting

Grafting experiments of H9-ePSC derived cortical progenitors (NPCs) were performed as described previously (Espuny-Camacho I *et al*., 2017)^2^. In brief, time-controlled mating’s were performed to obtain pregnant females. Five days before transplantation, frozen neural progenitor cells (NPCs) were thawed in the neural maintenance growth medium and supplemented with ROCK inhibitor (Y-27632; 10 mM Calbiochem, #688000). Cells were incubated in a humidified chamber at 37°C with 5% CO2, with replacement of growth media every 48h until transplantation. On the day of grafting, 30 minutes before enzymatic dissociation, Revitacel (Thermo; #A2644501) was added to the growth medium. Cells were dissociated with StemPro™ Accutase™ Cell Dissociation Reagent (Sigma, # A1110501). The viability of the cells was assessed by trypan blue staining. Finally, the cells were suspended at 100,000 cells/μl^−1^, or 50,000 cells/μl^−1^ concentration in Leibovitz’s L-15 Medium (Thermo; # 11415064) supplemented with 34 mM glucose solution.

### Grafting

Grafting was performed as previously described in Espuny-Camacho I et al., 2017^2^. In brief, grafting took place at the P1-P2 stage. Cryoanesthesia was used for grafting the NPCs. A small incision was made at the injection site (coordinates from bregma: anteroposterior, −1 mm; lateral, ±1 mm) using a sterile surgical blade. For unilateral grafting, 1 µl of the 100,000 cells/μl cell suspension, and for bilateral grafting 1 µl of 50,000 cells/μl was injected using 26G Hamilton syringe, without stereotactic frame support. After the injection, pups were allowed to recover under a heat lamp at 37 °C. Once fully recovered, they were placed back into the cage along with bedding material. Grafted pups were monitored daily for a week.

### Sample isolation

For immunohistochemistry of the brain, mice were anesthetized with an overdose of sodium pentobarbital, then perfused transcardially first with PBS followed by 4% PFA in PBS. Collected brains were postfixed overnight in 4% PFA. Brains were embedded in 4% Top Vision low-melting-point agarose (Thermo Fisher Scientific) and cut into 40 µm thick transverse coronal free-floating serial sections using a vibratome (Leica VT1000S). Brain sections were stored in cryoprotectant solution (30% ethylene glycol, 30% glycerol, 40% PBS) at −20 °C until use. For RNA extraction from grafts, animals were euthanized using cervical dislocation. The brain samples were quickly isolated and rinsed in ice-cold PBS containing RNasin (0.2u/ul). Then the brain was sliced into 1 mm thick sections using a cold brain matrix. Sections were collected immediately into cold PBS containing RNasin. These sections were placed under a fluorescent dissection microscope, the fine dissection of the GFP-positive areas was performed, and collected tissue was snap-frozen in liquid nitrogen. The rest of the brain was snap-frozen to be used for insoluble tau isolations. Samples were stored at -80°C until use. On the day of RNA isolation, samples were thawed on ice, and the Qiagen RNeasy mini kit was used for RNA purification. Blood samples were collected in EDTA-coated tubes during transcardial perfusion. The tubes were incubated on ice for 30 minutes, centrifuged at 2000g, and plasma supernatant was collected. Plasma samples were stored at -80 °C until use.

### Immunofluorescence and imaging

For immunofluorescence (IF) analysis, vibratome sections were washed three times with PBS to remove the residual storing solution. Where necessary, antigen retrieval was performed by microwave boiling in 10 mM tri-sodium citrate buffer pH 6.0. Then the sections were placed in permeabilization/blocking buffer containing 5% serum, corresponding to the host species of the secondary antibody (Donkey) in PBST (PBS with 0.20% Triton X-100 in 1xPBS) for one hour at room temperature. After the blocking step, the primary antibody (Supplementary Table 8) was added to the blocking solution and gently agitated overnight at 4°C. The next day, sections were washed three times for 5 minutes in PBST. Respective secondary antibodies were added, and sections were incubated for two hours at room temperature. If appropriate, DAPI staining was performed. Subsequently, the samples were washed three times for 5 minutes in PBST and mounted onto glass slides using the Glycergel mounting media and allowed to dry at room temperature. The slides were kept at 4°C until imaging. Confocal images were obtained using a Nikon Ti-E inverted microscope equipped with an A1R confocal unit driven by NIS (4.30) software. For excitation, 405 nm, 488 nm, 561 nm, 638 nm laser lines were used. All the images were obtained using a 20x (0.75 NA) objective lens, and Z-stack series of images of the area of interest were acquired using the NIS software. Roughly 10-12 Z-sections with 1 mm thickness were obtained. All images were acquired using similar acquisition parameters such as 16-bit, 1024x1024 quality, and images were processed in the FIJI/Image J software. All the images of Z-series stacks were then converted to Fiji/ImageJ maximum intensity projections. For quantification of percent positive cells, three to four sections comprising human xenograft were randomly selected and imaged using the above-described procedure and manually counted using NIS software.

### Quantification of the Immunofluorescence images

For quantification of the plaque-associated microglia and astroglia, a 20 µm ring was drawn around the plaque in the Nikon analysis software (NIS-Elements AR, Version 5.21.00). The number of microglia and astrocytes in the 20 µm ring were counted. A semi-automated house-made macro in NIS-Elements AR software was employed for quantifying the NP-tau in NIS-Elements AR software. In brief, X34-positive plaques were identified with threshold function, and the volume around the plaque was expanded by 20 µm in the z-stack images. AT8, PHF1, or MC1-positive volume (3D) within the 20 µm area was measured, normalized against the total expanded volume around the plaque, and represented as percent positive volume in the 20 µm X34 ring. N is > 100 plaques per mouse and at least 4 mice from each genotype were used for quantification. For quantifying pRIPK1 (n= 5-6), pRIPK3 (n= 5-6), and pMLKL (n= 5-6) positive neurons, random sections were taken and N>1000 HUNU-positive human neurons and neurons co-labeled with HUNU, and the above-selected markers were counted and represented as percent positive neurons.

### Gallyas silver staining

The Gallyas silver staining method was used to stain the tau tangles^3^. The 40 µm vibratome sections were mounted onto positively charged microscope slides (Thermo; #6776214). The sections were allowed to dry for 24 hours before staining. The next day, slides were washed in distilled water for one minute and transferred immediately to alkali silver iodide (4 g sodium hydroxide, 10 g potassium iodide, 3.5 mL of 1% silver nitrate in 100 mL distilled water) for 1 minute. Next, slides were washed 3 times for 1 minute in 0.5% acetic acid. Then the slides were incubated in physical developer solution (Solution 1: 50 mg sodium carbonate in 1 liter; Solution 2: 2 mg ammonium nitrate, 2 mg silver nitrate, 10 mg tungstosilicic acid in one-liter distilled water; Solution 3: 2 mg ammonium nitrate, 2 mg silver nitrate, 10 g tungstosilicic acid, 7.3 ml 35% formaldehyde in one-liter distilled water) for 15-30 minutes. The developer was made fresh before use in a 10:4:6 ratio (solution 1: solution 2: solution 3). Next, the samples were washed in 0.5% acetic acid and 1% sodium thiosulfate for 5 minutes each. Subsequently, samples were washed in distilled water for five minutes, incubated in 0.5% gold chloride solution, and sodium thiosulfate for 5 minutes each. Finally, the samples were washed in distilled water for five minutes, counterstained with nuclear fast red, and mounted using Permount solution (Fischer Scientific; #15820100).

### Insoluble tau isolation

Mouse brain samples were homogenized using nine volumes (v/w) of high-salt buffer (10 mM Tris-HCl, pH 7.4, 0.8 M NaCl, 1 mM EDTA, and 2 mM dithiothreitol [DTT], with protease inhibitor cocktail, a phosphatase inhibitor, and PMSF) with 0.1% sarkosyl and 10% sucrose and centrifuged at 10,000 g for 10 min at 4°C. The resulting pellets were reextracted twice using the same buffer, and the supernatants from all three extractions were pooled. Next, sarkosyl was added to the pooled supernatant to reach 1% final concentration. Samples were gently shacked for one hour at room temperature, followed by centrifugation at 300,000 g for 60 min at 4°C. The resulting pellet, which contains 1% sarkosyl-insoluble tau, was washed once with PBS, and resuspended in PBS (∼100µl/gm tissue) by passing through a 27G needle. Then the sample was subjected to a brief sonication (20 pulses at 1 s/pulse) using a water bath sonicator. Samples were centrifuged at 100,000 g for 30 min at 4°C to remove the major protein contaminants and to enrich tau filaments. The resulting pellet from this step is resuspended in PBS and further sonicated with 60 short pulses (1 s/pulse). Final low-speed centrifugation (10,000 g) was carried for 30 minutes at 4°C to remove contaminating large debris. Finally, enriched tau PHFs from the supernatant were separated and stored at -80°C until use.

### Negative staining and immuno-Electron Microscopy

For immuno-EM, diluted sarkosyl-insoluble fractions of the tau filaments were loaded onto the glow-discharged (Leica EM AC600) 400 mesh formvar/carbon film-coated copper grids (#01754-F; Ted Pella Inc) for 5 minutes and washed three times with PBS and blocked for 10 min with 5% acetylated BSA (Aurion). The grids were incubated with primary antibodies in a blocking buffer for three hours at room temperature followed by three times five minutes wash with PBS. Then the grids were incubated with 10 nm gold-conjugated secondary antibodies (1:20) in blocking buffer for 2-3 hours at room temperature. After that, grids were washed three times with sterile water and stained with 2% uranyl acetate for one minute. Finally, the grids were dried and imaged using a JEM-1400 transmission electron microscope (JEOL), equipped with an 11Mpixel Olympus SIS Quemesa camera.

### Cell number determination using qPCR

The number of human cells in the xenografted mice is determined as described previously^4–6^. In brief, total DNA from the xenografted mice is extracted using the TRI Reagent™ Solution (Thermo Fischer, #AM9738) according to the manufacturer’s instructions. Using human-specific primers, a standard curve has been established with human genomic DNA (gDNA) alone. The specificity of the primers was assessed by spiking an increasing amount of mouse gDNA. 80ng of DNA from the xenografted mice samples is used for the final qPCR assay. An absolute number of human cells was calculated by interpolating the standard curve and considering the standard reference of 6 pg of DNA/cell in the diploid human cell.

### RNA extraction from human brain

For qPCR analysis, frozen brain samples from the superior occipital gyrus or superior parietal gyrus of AD and age-matched control samples obtained from Netherlands Brain Bank were used (Supplementary Table 8). According to the manufacturer’s instructions, RNA was extracted from the brain samples using TRI Reagent™ Solution (Thermo Fischer, #AM9738).

### RNAScope

Cryostat sections from fresh frozen brain samples from the temporal gyrus (Supplementary Table 8) were used for the RNAScope based detection. RNAScope was performed according to the manufacturer’s instructions using a human *MEG3* -specific probe.

### Plasma P-tau181 measurements

Plasma samples were thawed at room temperature, vortexed for 30 seconds, and then centrifuged for 10 minutes at 4000 *g* at room temperature. Plasma P-tau181 concentration was measured at the Clinical Neurochemistry Laboratory, University of Gothenburg (Mölndal, Sweden) in one analytical run on a Single-molecule array (Simoa) HD-X instrument (Quanterix, Billerica, MA, USA). All experimental samples were run in singlicate after diluting the samples 1:2 in Tau 2.0 buffer (Quanterix, Billerica, MA). At the beginning and end of the assay plate, two internal quality control samples with concentrations of 11.1 pg/mL and 4.0 pg/mL were run in duplicate, with intra-plate variability of 2.2% and 1.5 %, respectively. AT270 mouse monoclonal antibody (Invitrogen, Waltham, MA, USA) specific for the threonine-181 phosphorylation site was coupled to paramagnetic beads (Quanterix) and used for capture. As a detector, the anti-tau mouse monoclonal antibody Tau12 (BioLegend, San Diego, CA, USA) was used, which binds the N-terminal on human tau protein. The detection antibody was conjugated to biotin (Thermo Fisher Scientific, Waltham, MA, USA) following the manufacturer’s recommendations. Full-length recombinant Tau 1-441 phosphorylated *in vitro* by glycogen synthase kinase 3β (GSK3β) (SignalChem, Vancouver, BC, Canada) was used as the calibrator.

### Protein concentration determination

The total protein concentration of the protein samples was determined with the standard Bradford protein assay.

### Immunoblotting

Approximately 25-30 µg of protein was used for immunoblotting. Samples were diluted with sterile water up to 12 µl, and 4 µl of NuPAGE^TM^ LDS Sample Buffer (4x) (Thermo Fisher Scientific), supplemented with 4% beta-mercaptoethanol was added. Samples were boiled for 10 min at 70°C. Denatured samples were loaded onto a 4-12 % Bis-Tris Gel (Invitrogen) and run at 170 V for 1 h in NuPAGE^TM^ MOPS SDS Running Buffer (Thermo Fisher Scientific). Proteins were transferred to a 0.2 µm nitrocellulose blotting membrane (GE Healthcare) for one hour at 25 V in cold transfer buffer (NuPAGE^TM^, Thermo Fisher Scientific) with 10 % MeOH. Ponceau staining was performed, and the membranes were blocked for one hour with 5 % milk in 0.1 % Tris buffered saline (TBS) Tween20 (TBS-T). The primary antibodies were added in blocking solution and incubated overnight at 4°C. The next day, the membranes were washed three times with TBS-T and incubated for one hour at room temperature with the appropriate secondary antibodies (Supplementary Table 8). Blots were developed by adding Western Lightning Plus-ECL Substrates (PerkinElmer) and imaged with an ImageQuant LAS4000 (GE Healthcare). For semi-quantitative analysis of western blots, AIDA Image Analyzer v5.0 was used.

### MEG3 expression

*MEG3*expression was achieved by cloning the cDNA of MEG3 transcript variant-1 under EF1α promoter in a lentiviral vector backbone (VectorBuilder). A lentiviral vector containing CAG: GFP was used as a control vector. For expression in neurons, NPCs were thawed on PLO (Sigma-Aldrich, # P4957) and laminin (Sigma-Aldrich, #L2020) coated plates. NPCs were treated with 10 µM DAPT (Sigma-Aldrich, #D5942) for two days to facilitate faster maturation. Neurons were transduced with either control or *MEG3* construct at ∼DIV68.

### Cell viability

Cell viability experiments were performed using the CellTiter-Glo® Luminescent cell viability assay, according to the manufacturer’s instructions. In brief, cells were seeded in a 96-well plate and transduced with either control or *MEG3* vector at MOI-10. On the day of the assay, conditioned media was removed, and fresh 100 ul media was added along with 100 µl of the CellTiter-Glo® reagent. Cells were incubated at room temperature for 30 minutes on a shaker before reading the luminescence signal on an Envison plate reader.

### RNA-Seq alignment and pre-processing

For xenografted samples, reads were aligned to a joint human plus mouse reference genome (GRCh38 and mm10, build 102) using STAR v2.7.3^7^, and then counted using Subread’s featureCounts v2.0.1^8^, in both cases using default parameters. Read count matrices for human and mouse were treated separately for downstream analysis. We filtered out genes with a mean read count < 5. We achieved means of 13,355,157 human reads per sample, and 162,882,734 mouse reads per sample. For in vitro *MEG3* overexpressing samples, reads were aligned to the human GRCh38 reference (build 102) and counted as above. After filtering low-count genes (as for xenografted samples), a mean read count of 20,770,413 reads per sample remained (Extended Data Fig. 12a). Raw counts were normalized using the R package edgeR v3.34, specifically a weighted trimmed mean of M values (TMM)^9^. Principal components analysis on the normalized count data was performed using the prcomp() function in R. Xenograft sample (SB295) appeared as an outlier in both mouse and human and was excluded from the analysis. In the human data, SB197 and SB198 also separated out from other 2M samples. These two samples showed strong upregulation of cell cycle markers relative to the other 2M samples. They were not excluded.

### Differential expression

Differential expression (DE) analysis was performed using edgeR’s generalized linear model, testing for differential expression using likelihood ratio tests. False Discovery Rate < 0.05 was considered significant. For *MEG3* expression data, we set differentiation run as a covariate, as it was shown to be a confounder in a PCA analysis (Extended Data Fig. 12b). DE results are available in Supplementary table 6. For visualizing the data, Bland–Altman plot as (MA) were used.

### Comparison of Rag2^-/-^ and *non*-Rag2^-/-^ mouse transcriptomes

We compared our *Rag2^-/-^/App^NL-G-F^* response in mice to a previous amyloid mouse model response^10^. We extracted differential expression results from APP^swe^/PS1^L66p^ versus controls at 10 months. We plotted these log fold changes against our *Rag2^-/-^/App^NL-G-F^* log fold changes at 6M, using only genes present in both datasets. A correlation was calculated between genes that were significantly differentially expressed in one or both studies.

### Functional Enrichment Analysis

DAVID (The Database for Annotation, Visualisation and Integrated Discovery) was used to perform functional enrichment^11, 12^. REVIGO^13^ was used for visualizing gene ontology terms in semantic space, setting the ‘species’ option to *homo sapiens*, and using their SimRel algorithm to cluster similar terms. Terms with FDR < 0.1 were considered significant unless otherwise indicated. Functional enrichment results are available in Supplementary Table 3 for transplanted neurons. Functional enrichment results for *MEG3* upregulated and downregulated genes are in Supplementary Table 7 for transplanted neurons

### Gene set Enrichment Analyses

GSEA was performed using the fgsea package in R (v1.18.0) with default parameters, using the Benjamini-Hochberg correction. We used the log fold changes from human xenografted neurons in amyloid mouse versus their controls, at each time point, as ranks. Genesets used in these analyses are described below. When comparing *MEG*3 overexpression data to xenografted neuron data, we selected the top 400 *MEG3* overexpression upregulated genes and the bottom 400 *MEG3* expression downregulated genes, based on log fold change, as our genesets.

### Previous gene sets

For the dedifferentiation analysis, genesets were taken from^14–19^. Genesets are available in Supplementary Table 4. Genesets related to comparison of transplanted neurons with bulk AD brain transcriptome, related to Extended data fig. 8a, b are provided in the Supplementary Table 5.

---

**Extended Data Figure 1:**
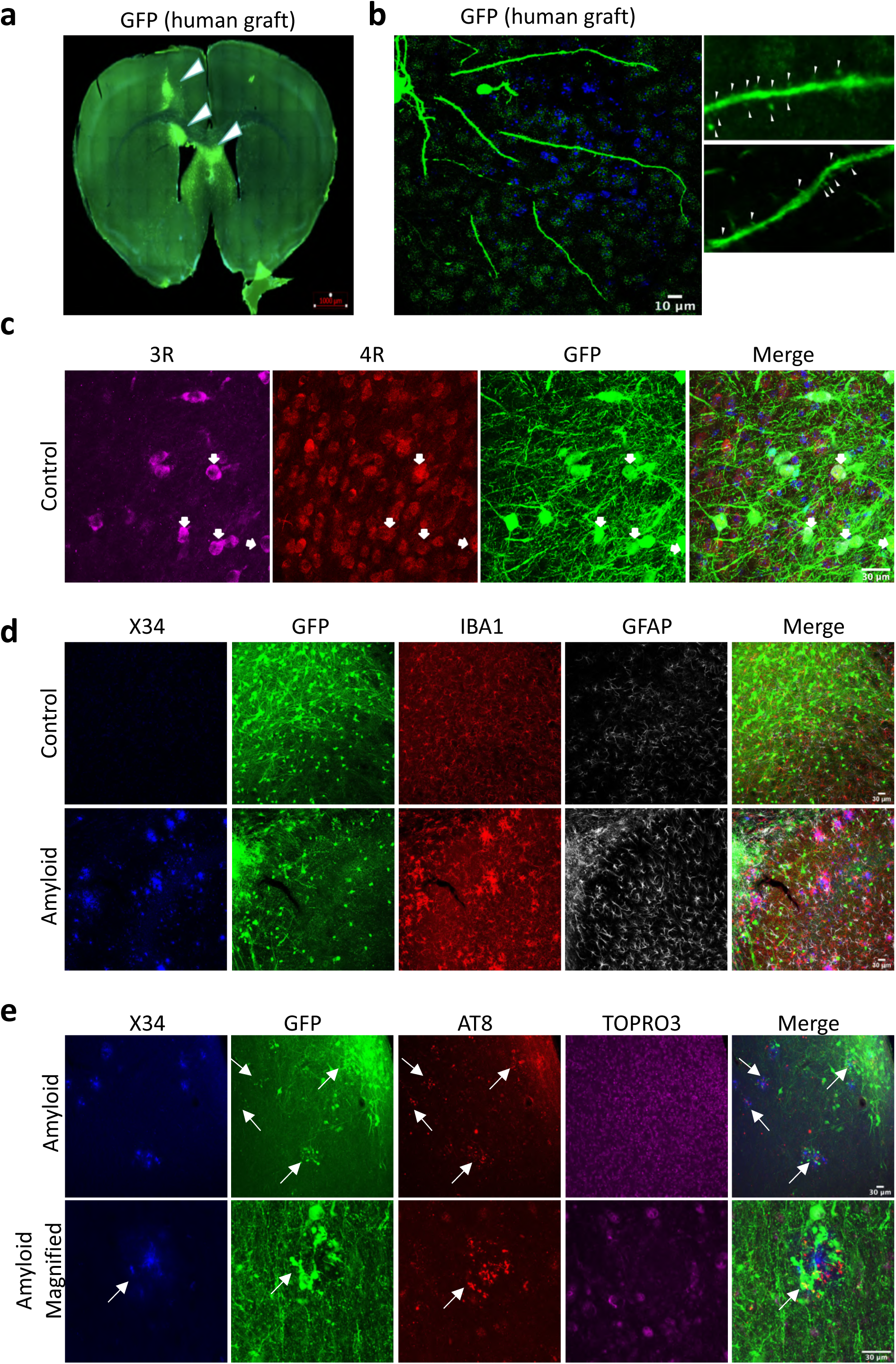
Integration of the transplanted neurons in the Rag2^-/-^ animals. **a,** Coronal section of 18-months old grafted mouse brain showing the graft location (green, indicated with white arrows). Scale bar 1000 µm. **b,** Higher magnification image of neuronal process displaying dendrites and mature spines (right boxes). White arrows indicates dendritic spines. Scale bar 10 µm. **c,** Representative confocal images of grafted neurons 6 months post-transplantation stained with 3R (magenta) and 4R (red) TAU isoforms (examples of human neurons indicated with white arrows). Scale bar 30 µm. **d,** Representative confocal images showing glial responses at 18 months post-transplantation. Amyloid (X34 in blue), human neurons in green (GFP), GFAP (grey), IBA1 (red). Scale bar 30 µm. **e,** Representative confocal images 6-months old grafted mice showing dystrophic neurites and neuritic plaque tau (NP-tau) pathology (AT8) around the X34 plaques 6 months post-transplantation. The top panel shows NP-tau, indicated with white arrows. The bottom panel shows a magnified region of one plaque showing dystrophic neurites and NP-tau. Amyloid (X34 in blue), human neurons in green (GFP), P-tau (AT8, red), TOPRO3 (nucleus, magenta). Scale bar 30 µm.

**Extended Data Figure 2:**
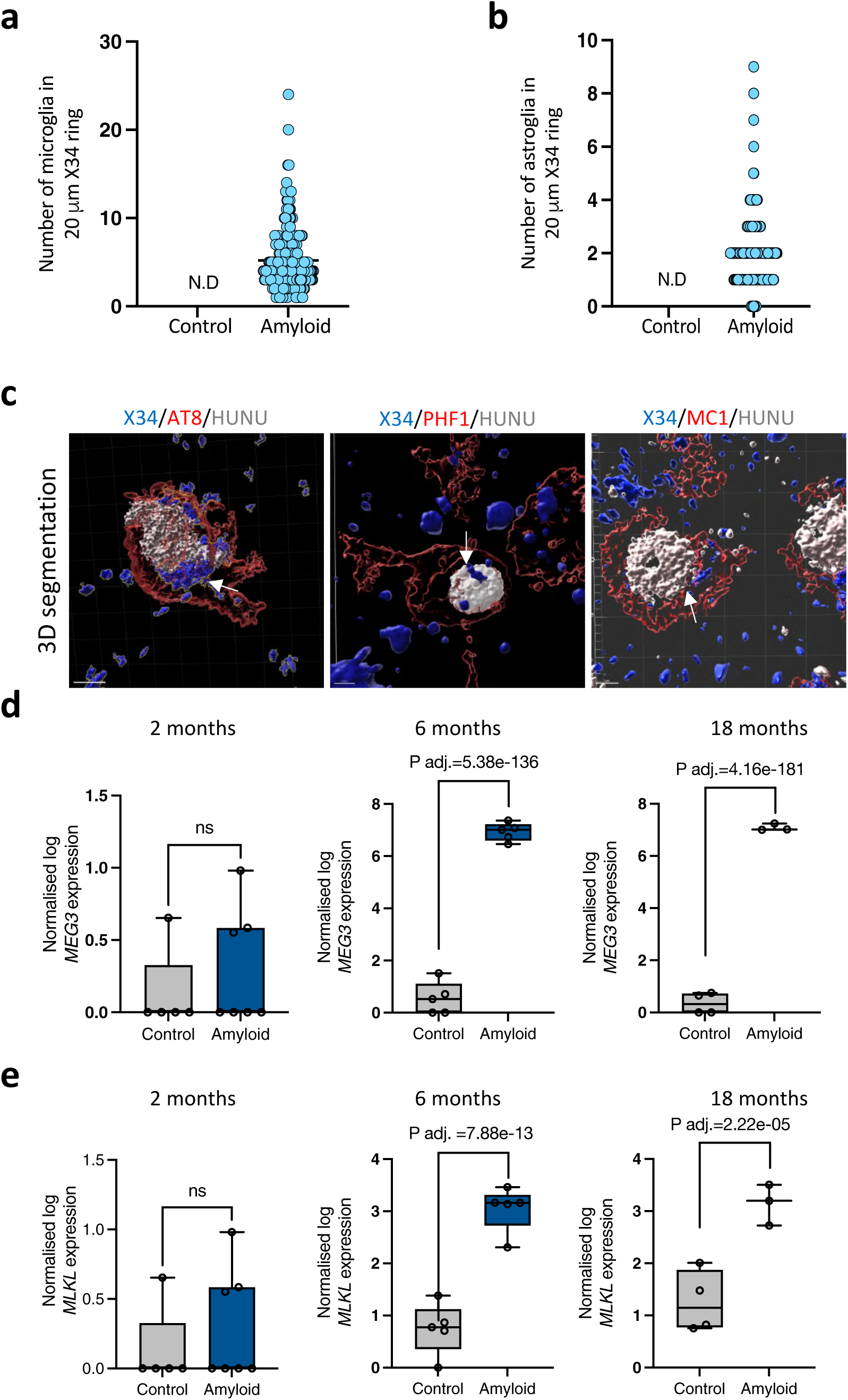
Characterization of the 18 months old grafted animals. **a,** Quantification of the number of host microglia around the Aβ plaque, within 20 microns diameter (n=4, >100 plaques/mice). **b,** Quantification of the number of host astrocytes around the Aβ plaque, within 20 microns diameter (n=4, >100 plaques/mice). **c,** Neurons immunoreactive to AT8, PHF1 or MC1 are segmented in Imaris software and rendered to reveal intracellular X34 staining (β-sheet fibrillary structures), an indicative of the intracellular tau fibrils. Extracellular X34 (blue) staining represents Aβ plaques where as intracellular X34 (blue) represents tau β-sheet structures. White arrows indicate X34 staining in neuronal somas. Amyloid (X34, blue), P-tau (red), human nucleus (HUNU, grey). Scale bar 5 µm. **d,** Human *MEG3* expression from 2-months (control n=5, amyloid n=7), 6-months (control n=5, amyloid n=5), and 18-months (control n=4, amyloid n=3) post-transplantation. **e,** Human *MLKL* expression from 2-months (control n=5, amyloid n=7), 6-months (control n=5, amyloid n=5), and 18-months (control n=4, amyloid n=3) post-transplantation.

**Extended Data Figure 3:**
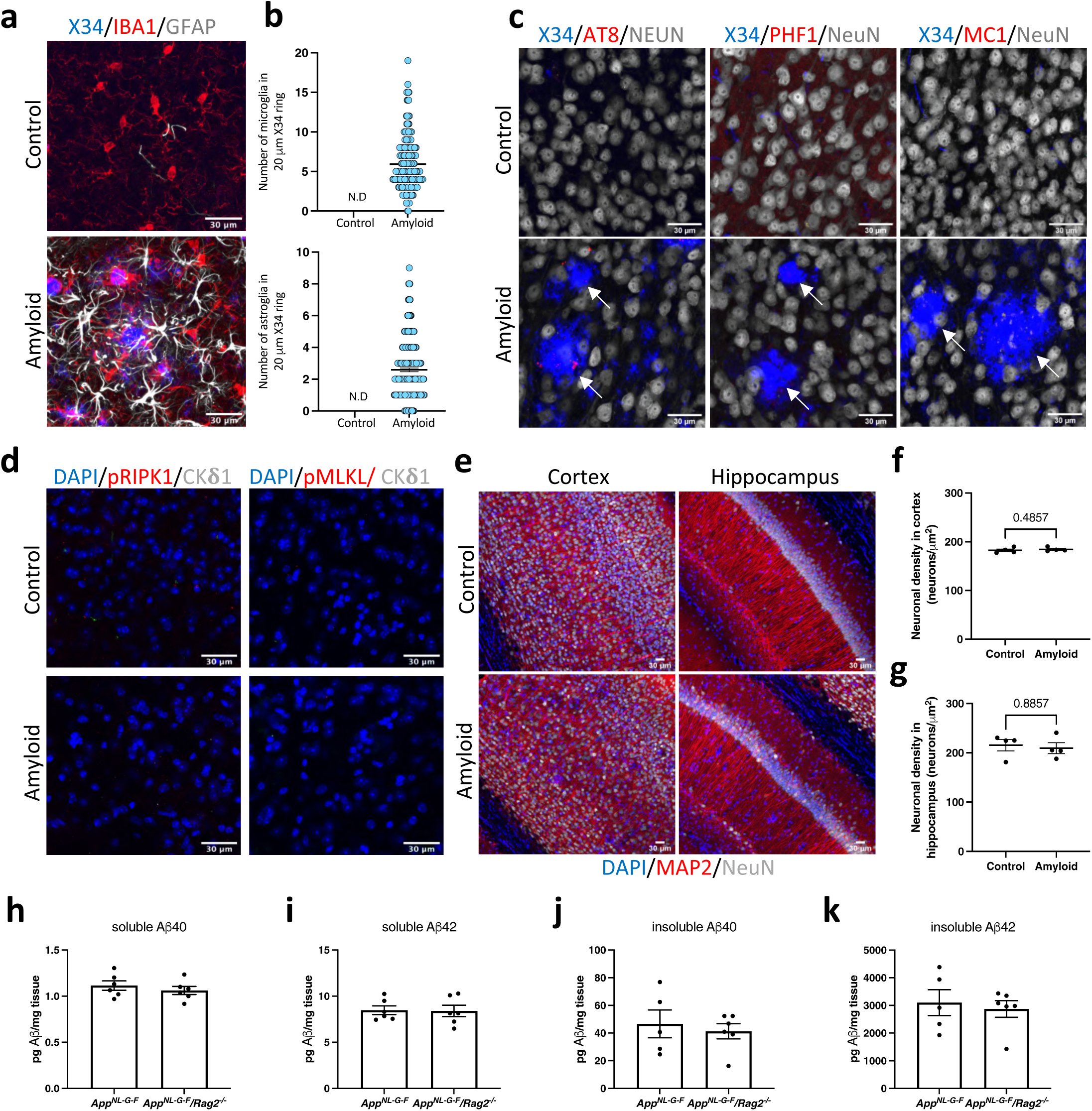
Characterization of 18 months old non-grafted animals. **a,** Representative confocal images of non-grafted 18 months old control (top) (n=4) and amyloid mice (bottom) (n=4) showing the glial reaction. Amyloid (X34, blue), microglia (IBA1, red), Astrocytes (GFAP, grey). Scale bar 30 µm. **b,** Quantification of the number of host microglia (top) and astroglia (bottom) around the Aβ plaque, within 20 microns diameter (n=4 mice, >100 plaques/mice). ND, not observed. **c,** Representative confocal images of neuritic plaque tau stained using AT8 (P-tau Ser202, Thr205), PHF1 (P-tau Ser396), and MC1 (pathological conformation) in 18 months old control (n=4) and amyloid (n=4) animals. Amyloid (X34, blue), P-tau (AT8 or PHF1, red), pathological conformation of tau (MC1, red), neurons (NeuN, grey). Scale bar 30 µm. **d,** Representative confocal images of activated necroptotic markers from 18-months old control (n=4) and amyloid (n=4) mice. Amyloid (X34 in blue), pRIPK1 and pMLKL (red), GVDs (CK1δ, grey). Scale bar 30 µM. **e,** Representative confocal images displaying neuronal density from cortex and hippocampus of the 18-months old control (n=4) and amyloid mice (n=4). Nucleus (DAPI, blue), MAP2 (red), neurons (NeuN, grey). Scale bar 30 µM. **f,** Quantitative representation of the neuronal density in cortex in 18-months old control (n=4) and amyloid (n=4) mice. **g,** Quantitative representation of the neuronal density in hippocampus in 18-months old control (n=4) and amyloid (n=4) mice. **h,** Soluble Aβ40 levels in hippocampus of 6-months old *App^NL-G-F^* (n=6) and *Rag2^-^*^/-^/*App^NL-G-F^* (n=6). **I,** Soluble Aβ42 levels in hippocampus of 6-months old *App^NL-G-F^* (n=6) and *Rag2^-^*^/-^/*App^NL-G-F^* (n=6). **j,** Guanidine extractable insoluble Aβ40 levels in hippocampus of 6-months old *App^NL-G-F^* (n=5) and *Rag2^-^*^/-^/*App^NL-G-F^* (n=6). **k,** Guanidine extractable insoluble Aβ42 levels in hippocampus of 6-months old *App^NL-G-F^* (n=5) and *Rag2^-^*^/-^/*App^NL-G-F^* (n=6).

**Extended Data Figure 4:**
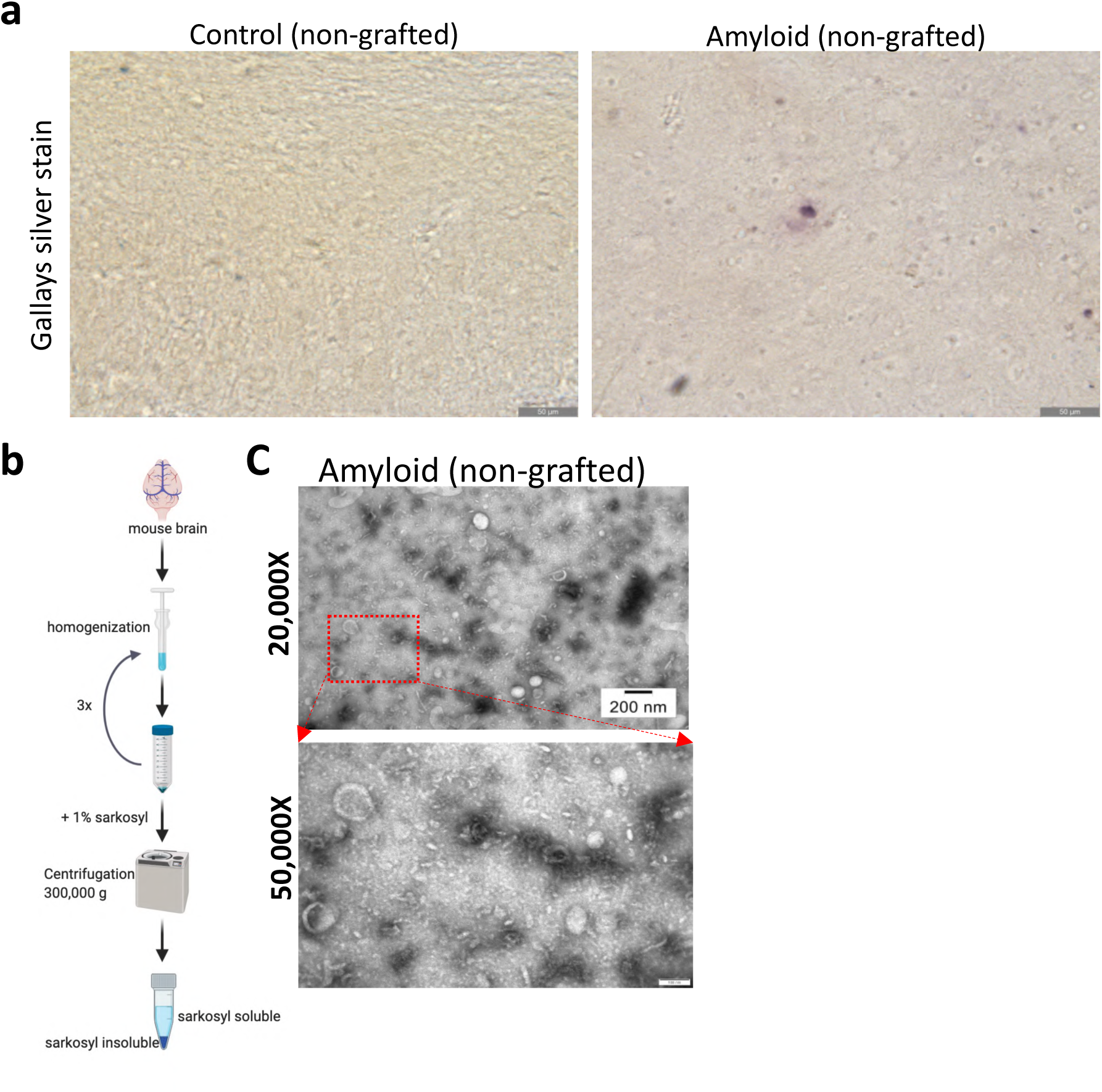
Characterization of TAU pathology in 18 months old non-grafted animals. **a,** Represenative light microscopic images of Gallays silver stain from 18-months old non-grafted control and amyloid animals. Scale bar 50 µm. **b,** Schematic representation of the sarkosyl-insoluble TAU preparation. **c,** Representative electron micrograph images of immunogold labelled sarkosyl insoluble fractions from 18-months old amyloid mice (n=4). Top panel indicates 20,000x magnification (scale bar, 200 nm), bottom panel indicates 50,000x magnification (scale bar, 100 nm).

**Extended Data Figure 5:**
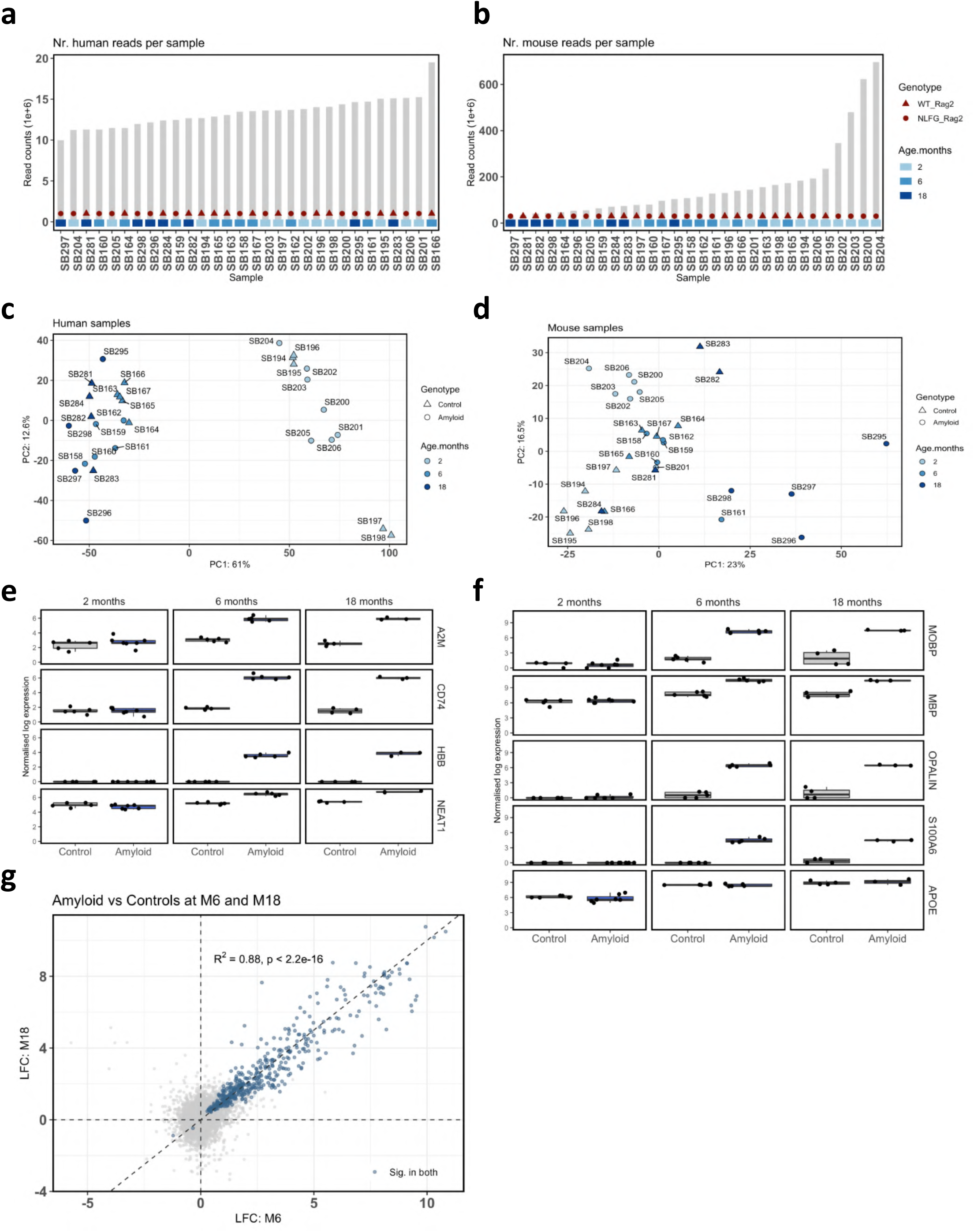
RNA sequencing of the human graft from 2-, 6– and 18-months post-transplantation. **a**, Representation of the total number of reads obtained from human graft in control and amyloid animals 2-months (control n=5, amyloid n=7), 6-months (control n=5, amyloid n=5), and 18-months (control n=4, amyloid n=4) post-transplantation. **b,** Representation of the total number of reads obtained from the host (mouse) tissue from 2-months (control n=5, amyloid n=7), 6-months (control n=5, amyloid n=5), and 18-months old (control n=4, amyloid n=3) animals. **c,** Principle component analysis (PCA) of the bulk RNA data from human grafts. **d,** PCA analysis of the bulk RNA data from host mouse tissue. **e,** Normalized gene expression levels of *A2M, CD74, HBB, NEAT1*, and *MOBP* in transplanted neurons at 2-, 6-, and 18-months post-transplantation. **f,** Log normalized gene expression levels of *MBP, OPALIN, S100A6,* and *APOE* in transplanted neurons at 2-, 6-, and 18-months post-transplantation. **g,** Scatter plot showing comparison of log fold changes (LFC) in human neurons between 6M and 18M. Genes highlighted blue are significant in both timepoints. The Pearson correlation (R^2^) of genes significantly changing in both models is 0.88.

**Extended Data Figure 6:**
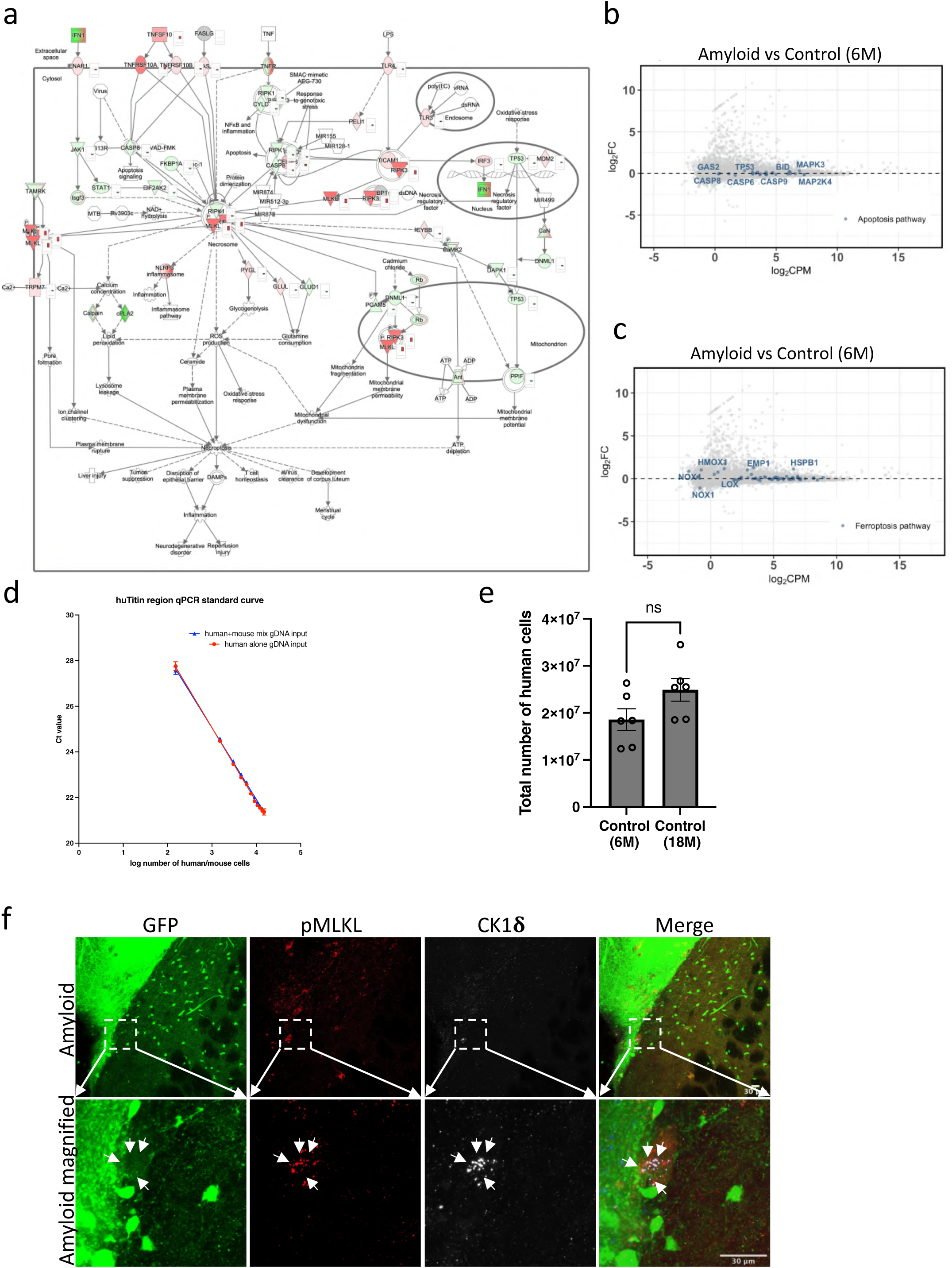
Neuronal cell death analysis in transplanted human neurons. **a,** Ingenuity pathway analysis (IPA) of canonical necroptosis pathway overlaid with differentially expressed gene from the 6-months old neuronal xenografts. Red indicates upregulation, and green indicates downregulation. **b,** Bland-Altman MA plot showing expression of apoptosis pathway genes in the 6 month old neuronal xenografts. **c,** Bland-Altman MA plot showing expression of ferroptosis pathway genes in the 6 month old neuronal xenografts. **d,** Establishment of the standard curve for qPCR using *TITIN* with human genomic DNA (red line) and human plus mouse genomic DNA (blue line). **e,** Quantitative representation of the human neurons in control mice (6-months and 18-months) (n=6). **f,** Representative confocal images showing co-localization of pMLKL with granulovacuolar degeneration (GVD) marker casein kinase 1 delta (CK1δ) in 18-months old amyloid mice. Human graft (GFP in green), pMLKL (red), CK1δ (grey). Scale bar 30 µM.

**Extended Data Figure 7:**
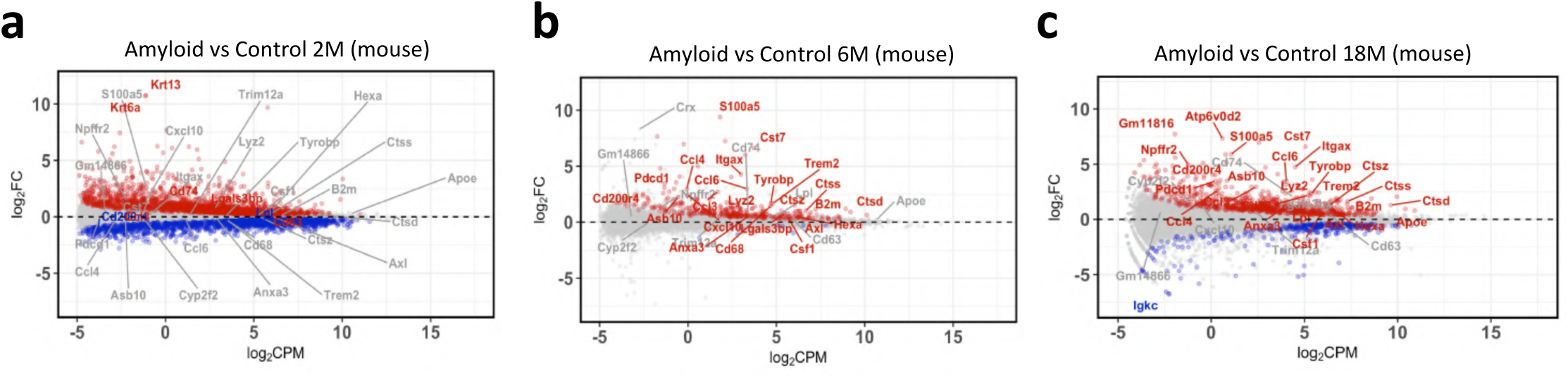
Gene expression profiles of mouse host tissue from mice at 2, 6 and 18 months. **a,** Bland-Altman MA plot showing the differential gene expression of mouse transcriptome at 2-months (control n=5, amyloid n=7), **b,** 6-months (control n=5, amyloid n=5), and **c,** 18-months (control n=4, amyloid n=3) old mice. Red, significantly upregulated genes. Blue, significantly downregulated genes (FDR < 0.05). Selected microglial activation genes are labelled. FC=fold change, CPM=counts per million.

**Extended Data Figure 8:**
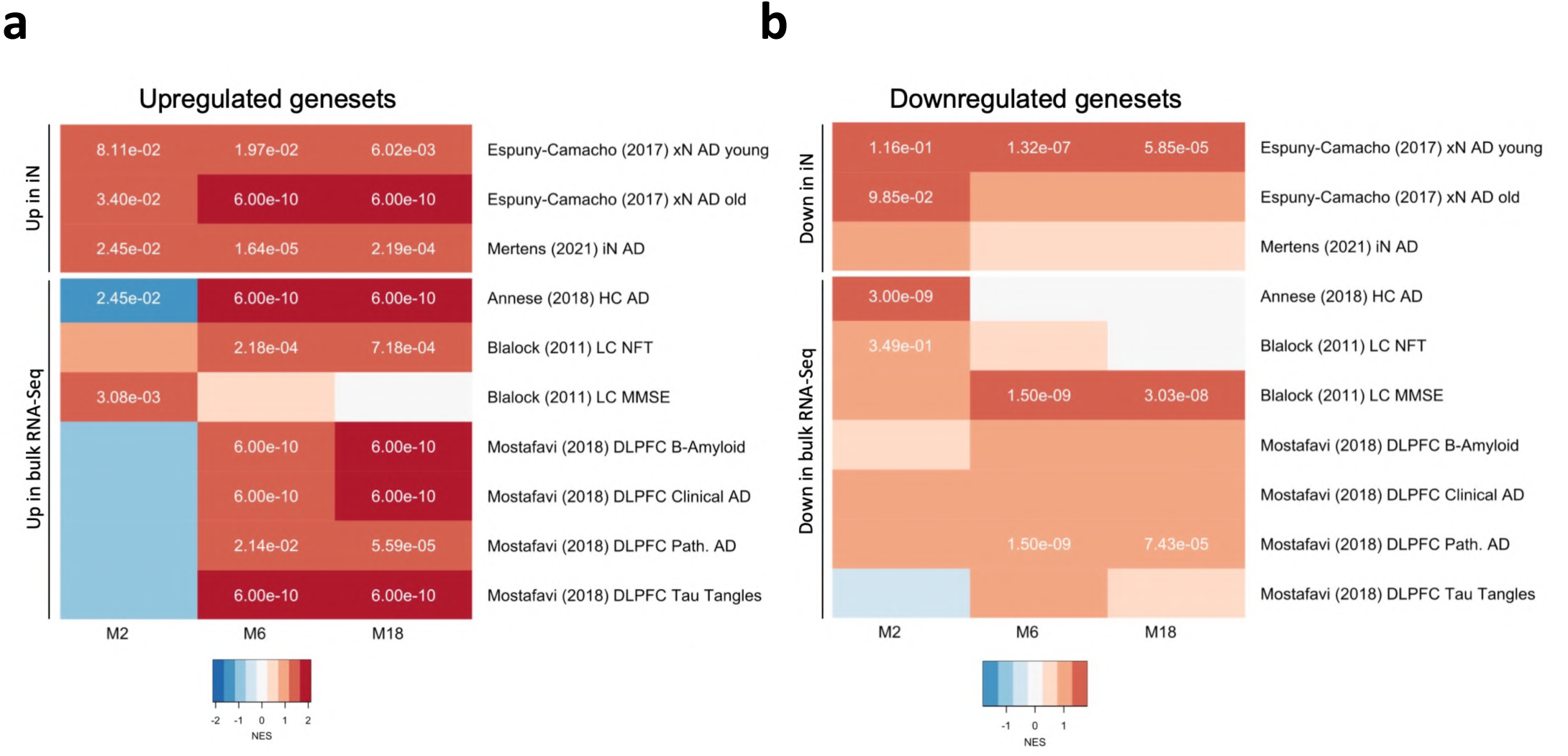
Xenografted neurons recapitulate signatures identified in human post-mortem tissue and other xenografted neurons. **a,** Heatmap of normalized enrichment scores (NES) in indicated gene sets ranked along the human graft fold changes (upregulated genes, amyloid vs control) shown in figure 2a-c. Positive enrichments are shown in red, negative enrichments are shown in blue. Significant FDR values (p.adj < 0.01) are shown as text in the boxes. M2, upregulated genes in human neurons 2 months post-transplantation (PT). M6, upregulated genes in human neurons 6M PT. M18, upregulated genes in human neurons 18M PT. **b,** Heatmap of normalised enrichment scores (NES) in indicated gene sets ranked along the human graft fold changes (downregulated genes, amyloid vs control) shown in figure 2a-c. Positive enrichments are shown in red, negative enrichments are shown in blue. Significant FDR values (p.adj < 0.05) are shown as text. M2, downregulated genes in human neurons 2 months post-transplantation (PT). M6, downregulated genes in human neurons 6M PT. M18, downregulated genes in human neurons 18M PT.

## Datasets used in the above comparison

Espuny-Camacho et al., 2017 xN AD young, DE upregulated genes form human neurons 4 months post-transplantation (PT). Espuny-Camacho et al., 2017 xN AD old, 6M-8M PT. (PMID: **28238547**).

Martens et al., 2021 iN AD, DE genes from neurons directly reprogrammed from the fibroblast-derived from AD patients compared to neurons reprogrammed from control fibroblasts. (PMID: **33910058**).

Annese (2018) HC AD, DE genes from Late-Onset AD (LOAD) compared to controls from the hippocampus. (PMID: **29523845**)

Blalock (2011) LC NFT, DE genes from laser captured P-tau positive neurons. (PMID: **21756998**) Blalock (2004) MMSE, DE genes from MiniMental Status Examination (MMSE>20) from AD brains. (PMID: **14769913**).

Mostafavi 2018 DLPFC, DE genes from dorsolateral prefrontal cortex from AD (PMID**: 29802388).**

**Extended Data Figure 9:**
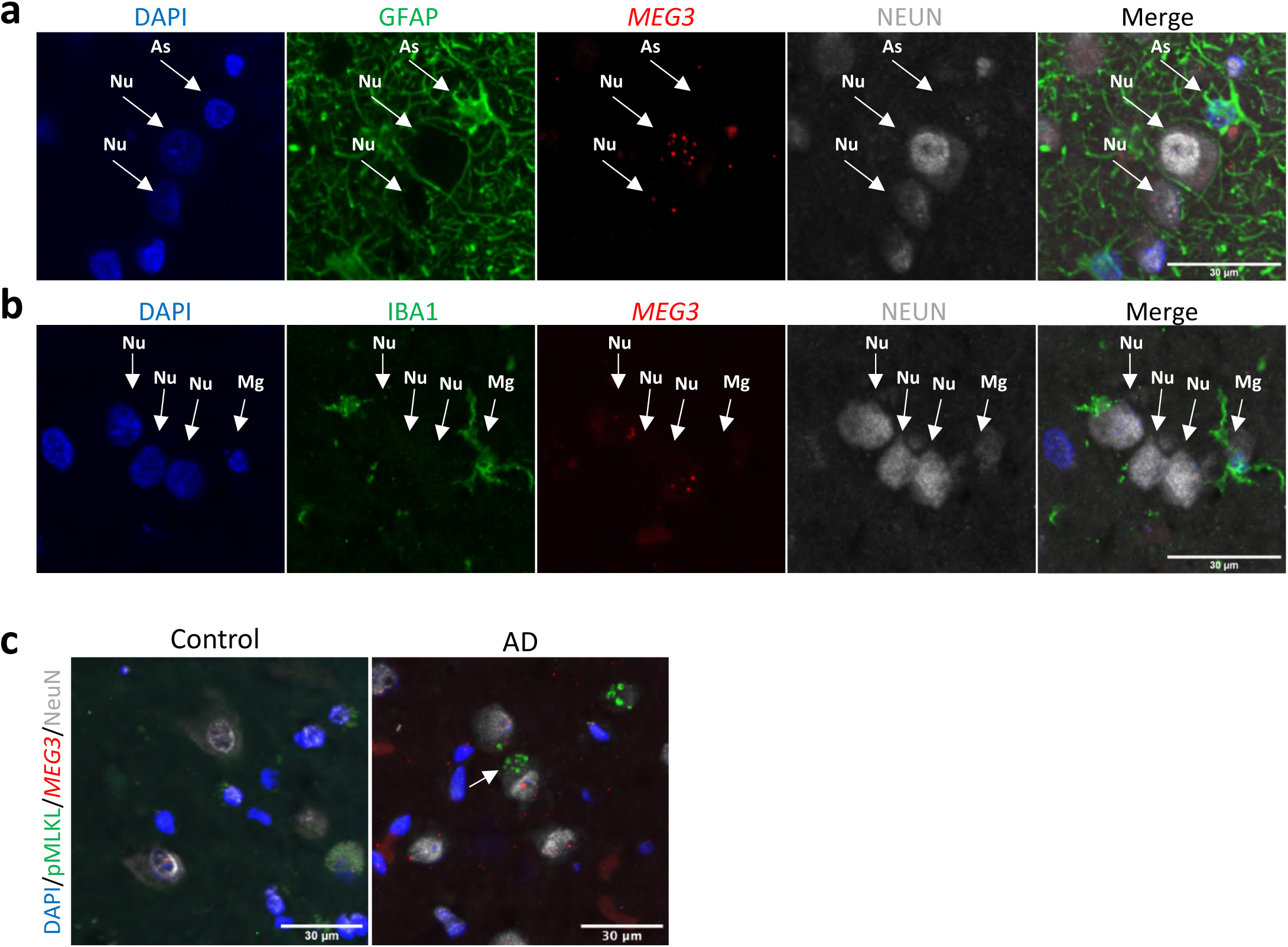
*MEG3* is selectively expressed in neurons. **a,** Representative confocal images from FFPE fixed human hippocampal brain samples stained with *MEG3* (RNA scope, red) and combined with immunostaining for GFAP (green), NeuN (grey), and DAPI (blue). Nu=Neuron, As=Astrocytes. Scale bar 30 µm. **b,** FFPE fixed human hippocampal brain samples were used for *MEG3* RNA scope (red) and combined with immunostaining for IBA1 (green), NeuN (grey), and DAPI (blue). Nu=Neuron, Mg=microglia. Scale bar 30 µm. **c,** Representative confocal images showing the colocalization of pMLKL (green) in the *MEG3* positive neurons in AD (n=3) and control (n=3) hippocampal brain samples. Scale bar 30 µm.

**Extended Data Figure 10:**
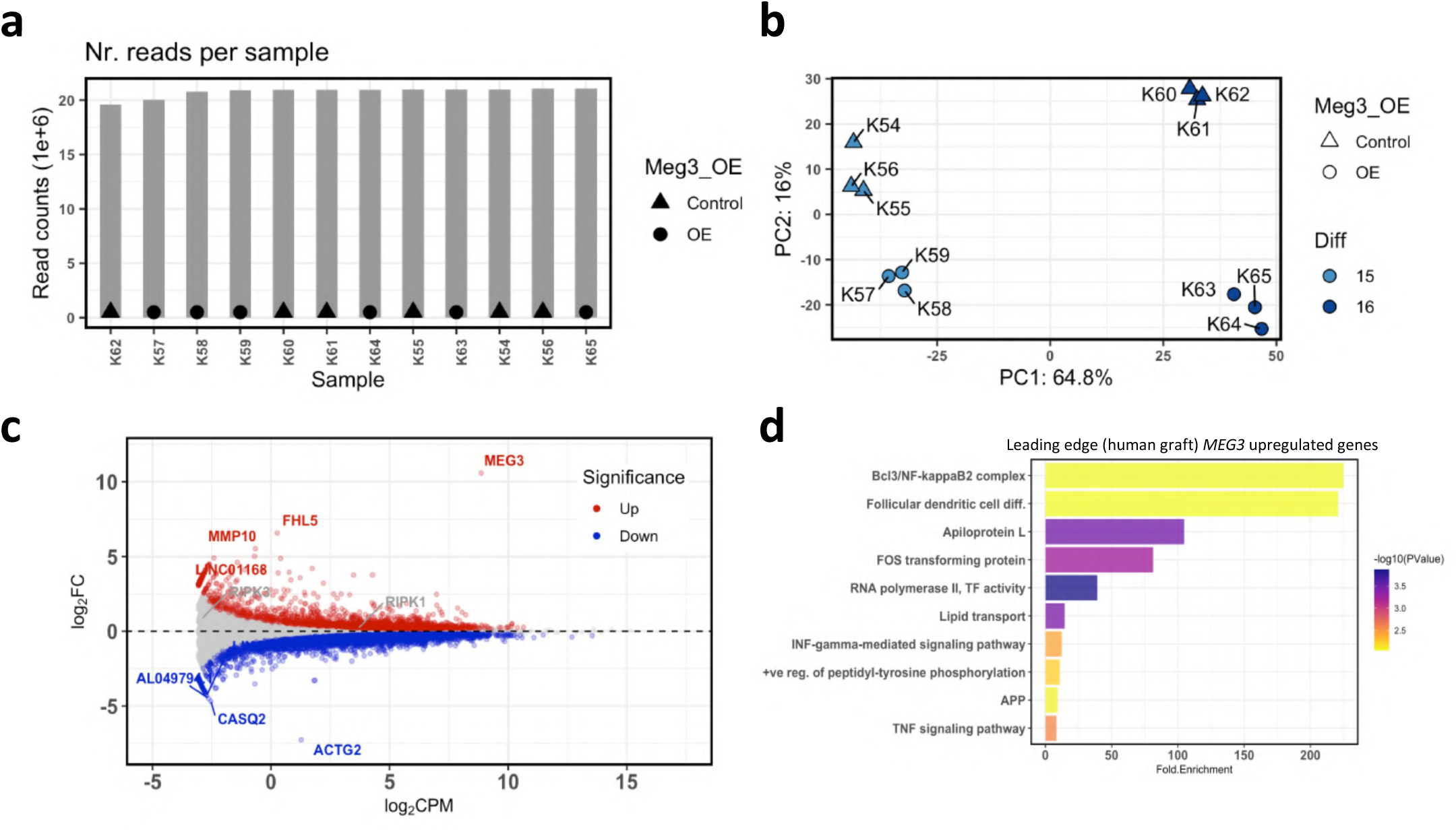
RNA sequencing of the *MEG3* expressing neurons. **a,** Barplots of the total number of reads per sample obtained from RNA sequencing of *MEG3* oexpression (OE, control n=6, overexpression n=6). **b,** Principle component analysis (PCA) of the RNA sequencing samples of *MEG3* expression (OE). Neurons derived from two independent differentiations (Diff), control LV (n=6), *MEG3* LV (n=6). **c,** Bland-Altman MA plot showing differential expression of bulk RNA sequencing of *MEG3* expression (control LC n=6, *MEG3 LV* n=6). Significantly upregulated genes are shown in red and downregulated genes in blue (FDR < 0.05). Full differential expression results are shown in Supplementary Table 6. **d,** DAVID gene ontology analysis of leading-edge upregulated genes from the GSEA analysis (indicated with red color in fig.3h). Top 10 GO terms were displayed. -ve=negative, +ve=positive, reg.=regulation , TF=transcription factor, TNF=tumor necrosis factor, APP=antigen process and presentation.

